# Availability of Zinc Impacts Interactions Between *Streptococcus sanguinis* and *Pseudomonas aeruginosa* in Co-culture

**DOI:** 10.1101/789099

**Authors:** Kewei Li, Alex Gifford, Thomas Hampton, George A. O’Toole

## Abstract

Airway infections associated with cystic fibrosis (CF) are polymicrobial. We reported previously that clinical isolates of *P. aeruginosa* promote the growth of a variety of streptococcal species. To explore the mechanistic basis of this interaction, we performed a genetic screen to identify mutants of *Streptococcus sanginuis* SK36 whose growth was no longer enhanced by *P. aeruginosa* PAO1. Mutations in zinc uptake systems of *S. sanginuis* SK36 reduced growth of these strains by 1-3 log compared to wild-type *S. sanginuis* SK36 when grown in coculture with *P. aeruginosa* PA01, while exogenous zinc (0.1-10 μm) rescued the coculture defect of zinc uptake mutants of *S. sanginuis* SK36. Zinc uptake mutants of *S. sanginuis* SK36 had no obvious growth defect in monoculture. Consistent with a competition for zinc driving coculture dynamics, *S. sanginuis* SK36 grown in coculture with *P. aeruginosa* showed increased expression of zinc uptake genes compared to *S. sanginuis* grown alone. Strains of *P. aeruginosa* PAO1 defective in zinc transport also supported more robust growth by *S. sanginuis* compared to coculture with wild-type *P. aeruginosa* PAO1. An analysis of 118 CF sputum samples revealed that total zinc levels varied from ~5-145 μM. At relatively low zinc levels, *Pseudomonas* and *Streptococcus* were found in approximately equal abundance; at higher zinc levels, we observed an increasing relative abundance of *Pseudomonas* and decline of *Streptococcus*, perhaps as a result of increasing zinc toxicity. Together, our data indicate that the relative abundance of these microbes in the CF airway may be impacted by zinc levels.

**IMPORTANCE:** Polymicrobial infections in CF likely impact patient health, but the mechanism(s) underlying such interactions are poorly understood. Here we show that interactions between *Pseudomonas* and *Streptococcus* are modulated by zinc availability using an *in vitro* model system, and clinical data are consistent with this model. Together with previous studies, our work supports a role for metal homeostasis as a key factor driving microbial interactions.

## INTRODUCTION

Cystic fibrosis (CF) is a monogenic autosomal recessive disorder caused by mutations in the cystic fibrosis transmembrane conductance regulator (CFTR) gene (1). It is estimated that ~70,000 individuals in the world are affected by CF and the most common mutation, caused by a deletion of phenylalanine at the 508th amino acid within the CFTR protein (ΔF508), is found in approximately 70% of this population (2, 3). CFTR dysfunction affects several body systems, and progressive lung disease due to chronic and recurrent microbial infections is the leading cause of morbidity and mortality in individuals with CF (4, 5). It has been shown that CF airway infections are polymicrobial (6, 7), and the composition and interspecies interactions within the polymicrobial communities can have profound and diverse consequences, including on bacterial growth (8–10), as well as disease progression and therapeutic outcomes (6, 11).

An example of such microbial interactions includes the CF-associated streptococcal species, the presence of which may influence the growth and/or virulence of other CF pathogens, including the important pathogen *Pseudomonas aeruginosa* (12, 13). In turn, *P. aeruginosa* can impact the growth and/or persistence of streptococci (8–10, 14), with the net impact of these interactions resulting in exacerbation (4, 15–17) or less server loss of lung function (11, 16, 18–21). Therefore, understanding how these pathogens interact with each other and their multicellular host to impact disease progression, as well as how these interactions are modified by the CF airway environment, is of high significance.

Zinc is an essential micronutrient for all organisms and serves as a structural or catalytic cofactor in 5–6% of proteins in bacterial proteome (22–24). However, high concentrations of zinc are toxic, possibly due to competition with other relevant metal ions (24, 25), inhibition of key enzymes and essential metabolic reactions in the cell (26–28) and inducing membrane stress (29). For most pathogens, there is intense competition for zinc in the face of the host’s defense mechanisms, called nutritional immunity (30), and conversely elevating zinc level is also a strategy used by macrophages to kill pathogens encapsulated in their phagosomes (31, 32).

To maintain zinc homeostasis, bacterial species have evolved multiple systems to import and export zinc in environments of zinc limitation or excess, respectively (33, 34). In *P. aeruginosa*, the two best-studied zinc acquisition systems are the high-affinity ZnuABC system (35–37) and the pseudopaline system (38, 39). ZnuABC is a member of the ATP-binding cassette (ABC) transporters, consisting of a zinc-specific binding periplasmic protein (ZnuA), an inner membrane permease (ZnuB), and an ATPase (ZnuC) (35, 37, 40). Loss of ZnuA in *P. aeruginosa* PAO1 results in an ~60% reduction in cellular zinc accumulation (35). The pseudopaline system is primarily involved in zinc uptake in zinc poor environments and zinc transport into the cell is achieved via the action of a four-gene operon (*cntOLMI*, also termed *zrmABCD*) (38, 39). This operon includes the genes for a TonB-dependent outer membrane pseudopaline receptor (CntO), two biosynthetic enzymes (CntL and CntM) responsible for synthesizing pseudopaline and an inner membrane transporter (CntI) involved in pseudopaline secretion (36, 38, 39). Both the ZnuABC system and the pseudopaline operon are negatively regulated by zinc level through the Fur-like zinc uptake regulator Zur (35, 37, 39), which can sense and respond to femtomolar changes of cytosolic zinc concentrations (41). When zinc is bound, Zur represses transcription of the zinc uptake systems and loss of Zur results in higher cytoplasmic zinc in *P. aeruginosa* (36, 37).

Similarly, several zinc transporters have been identified to contribute to zinc homeostasis and virulence in pathogenic streptococci (42, 43). In most streptococci, zinc acquisition involves a high-affinity zinc-ABC transporter, which is comprised of an integral membrane component (AdcB), an ATPase (AdcC) and one or several zinc-binding proteins (AdcA, AdcAII, Lbp, or Lmb) (42, 44–46). The streptococcal AdcR repressor, a MarR family regulator, is involved in regulation of this transporter (47).

Recent studies have suggested that total zinc levels are much higher in sputum from CF patients compared with healthy controls (48), but zinc availability is limited in the lung mucosa, leading to zinc starvation for *P. aeruginosa* during CF lung infection (49). Moreover, the host also employs zinc starvation or intoxication to retard streptococcal growth during colonization and infection (42, 43, 50). Nevertheless, the potential significance of zinc in the interplay between *P. aeruginosa* and *Streptococcus*, especially in the context of a polymicrobial infection in the CF airway, remains to be explored.

We recently showed that *P. aeruginosa* can enhance the growth of multiple oral *Streptococcus* spp. in coculture conditions (8, 10), but the molecular mechanism underlying such interactions is still poorly understood. In the present work, we used a comprehensive *Streptococcus sanguinis* SK36 mutant library to determine the genetic requirements for *S. sanguinis* SK36 to benefit from interaction with *P. aeruginosa*. Our results show that efficient zinc acquisition in *S. sanguinis* SK36 plays a critical role in *P. aeruginosa*-induced growth enhancement. We observed increased transcription of zinc uptake genes in *S. sanguinis* SK36 and demonstrate that *P. aeruginosa* and *S. sanguinis* SK36 are competing for zinc during co-cultivation. Furthermore, by coupling analysis of microbial communities and zinc content within CF sputum samples, we discovered a relationship between the relative abundance of *Streptococcus* and *Pseudomonas* with the concentrations of total zinc in CF sputum. These results suggest that zinc availability may play a role in *Pseudomonas*-*Streptococcus* interactions in vivo, and furthermore, changes in zinc levels may also be related to the development of the respiratory microflora in patients with CF.

## RESULTS

### *P. aeruginosa* and its conditioned medium stimulate *S. sanguinis* growth on both plastic and CF-derived airway cells

We have previously shown that *P. aeruginosa* can enhance *S. sanguinis* SK36 growth either as planktonic or biofilm cells in coculture compared to that of *S. sanguinis* SK36 monoculture, while the *P. aeruginosa* population was not significantly impacted by the presence of *S. sanguinis* (8, 10), a finding confirmed here Fig. S1A. Furthermore, an assay monitoring biofilm formation over time shows that the higher viable count of *S. sanguinis* SK36 in coculture with *P. aeruginosa* is due to enhanced growth of *S. sanguinis* SK36 after a modest reduction in viability (Fig. 1A). Interestingly, *S. sanguinis* SK36 did show enhanced growth at later time points in monoculture, indicating that *P. aeruginosa* may stimulate transition out of lag phase as one possible mechanism of the observed increased viable count of *S. sanguinis* SK36 observed in the coculture assay system. The viable counts of *P. aeruginosa* were similar in monoculture and coculture at all timepoints (Fig. 1B).

**Figure 1.**
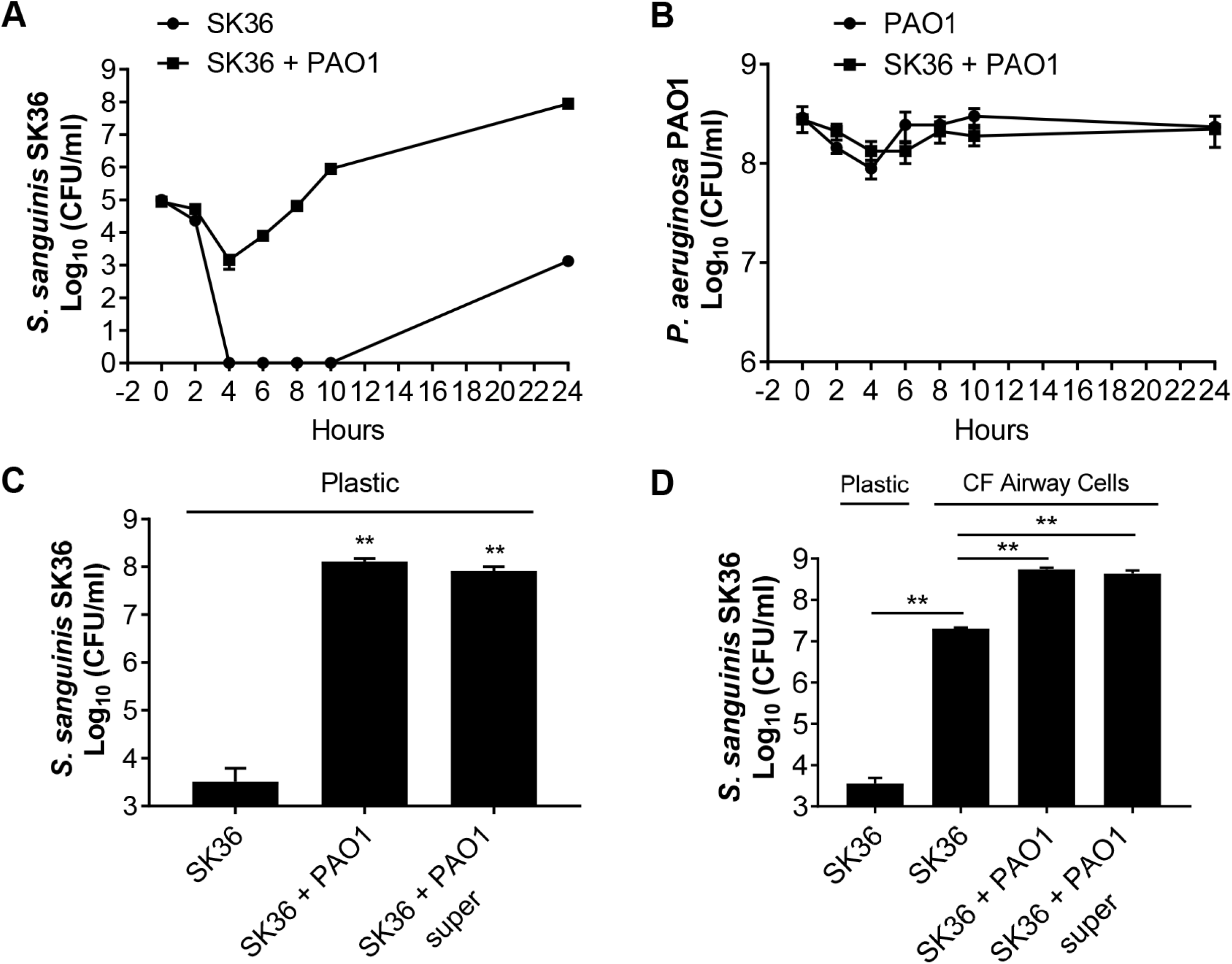
*P. aeruginosa* can stimulate the growth of *S. sanguinis* when grown in co-cultures. (A and B) Growth kinetics of *S. sanguinis* SK36 (labeled SK36) and *P. aeruginosa* PAO1 (labeled PAO1) in a 96-well deep well plate either in coculture or as a monoculture. (C-D) *P. aeruginosa* cells or a 1:2 dilution of *P. aeruginosa* supernatant (super) enhances *S. sanguinis* growth in coculture on a plastic substratum (C) or CFBE monolayers (D). Error bars indicate standard deviation of the means from a representative triplicate assay. (**, *P* < 0.01, Student’s *t*-test).

In an attempt to understand the basis of this observed growth enhancement of *S. sanguinis* SK36, we tested whether a cell free supernatant of *P. aeruginosa* PAO1 could stimulate the growth of *S. sanguinis* SK36. We found that *P. aeruginosa* cell-free supernatant was indeed capable of promoting *S. sanguinis* biofilm growth on plastic (Fig. 1C). We noted a significant increase in the number of *S. sanguinis* cells grown in the presence of a 1/2 or 1/4 dilution of the original *P. aeruginosa* PAO1 supernatant (Fig. S1B), with a dose-dependent decrease in growth enhancement with increasing dilution of the supernatant. Undiluted *P. aeruginosa* PAO1 supernatant showed a less robust effect on *S. sanguinis* SK36 growth than the 1/2 or 1/4 dilution, perhaps due to the known antimicrobial factors in *P. aeruginosa* supernatant, including siderophores, phenazines, cyanide and elastase (51–53). Taken together, these data suggest that one or more secreted factors in the *P. aeruginosa* PAO1 supernatant can enhance growth of *S. sanguinis* SK36.

To determine whether the interactions between *Streptococcus* and *P. aeruginosa* also occur when these microbes are grown in the presence of airway cells derived from patients with CF, we extended our observation to CF-derived bronchial epithelial cells homozygous for the ΔF508 allele of CFTR (referred to as “CFBE monolayers” hereafter) (54). We evaluated *S. sanguinis* SK36 growth on CFBE monolayers in the absence or presence of *P. aeruginosa* cells or supernatants. As a control, *S. sanguinis* growth on tissue culture plates without host cells was also included (Fig. 1D, left-most column, plastic). Interestingly, *S. sanguinis* growth enhancement was observed with monocultures biofilms on host cells, and *P. aeruginosa* cells and supernatants further enhanced the growth of *S. sanguinis* SK36 (Fig. 1D). Collectively, our results demonstrate that the ability of *P. aeruginosa* PAO1 cells and supernatant to promote the growth of *S. sanguinis* SK36 under a variety of conditions.

### Identification of *S. sanguinis* mutants defective in growth enhancement mediated by *P. aeruginosa*

To identify *S. sanguinis* genes that are potentially required for the increase in *S. sanguinis* SK36 growth in the *P. aeruginosa* PAO1-*S. sanguinis* SK36 coculture, we performed a genome-wide screen of an available comprehensive *S. sanguinis* SK36 mutant library (55), which covers ~90% of the predicted 2270 protein coding genes in the *S. sanguinis* SK36 genome. In this screen we sought to identify mutants with reduced enhancement phenotypes in the presence of *P. aeruginosa* PAO1 (Fig. 2A). Of 2048 mutants screened, a total of 80 mutants showed a measurable and repeatable reduction in the enhancement of *S. sanguinis* growth (Table S1).

**Figure 2.**
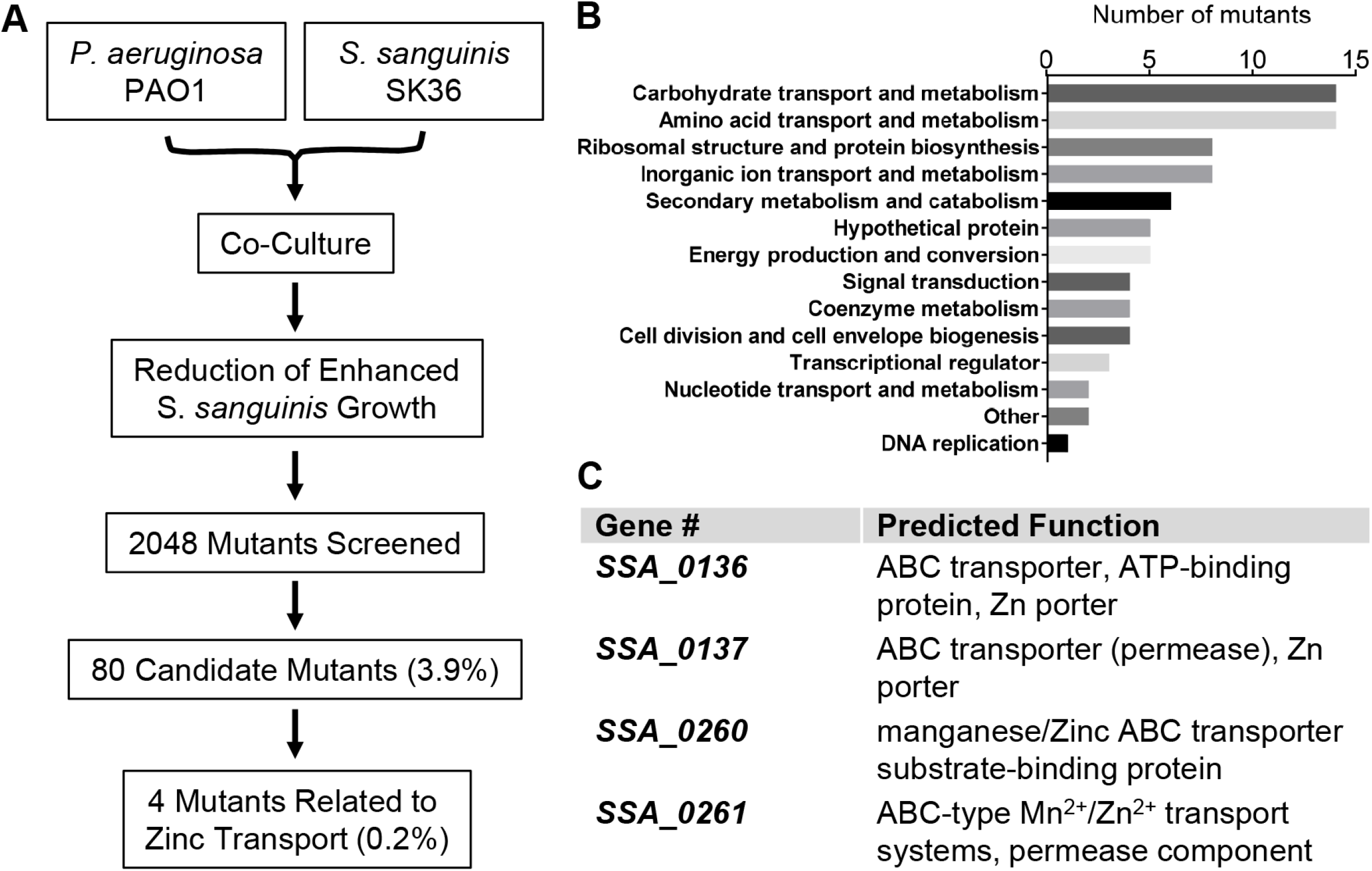
Identification of *S. sanguinis* SK36 mutants that exhibits reduced growth enhancement mediated by *P. aeruginosa*. (A) Schematic diagram of the genome-wide screen of *S. sanguinis* SK36 mutant library. (B) Functional classification of screen hits using KEGG Orthology (KO) database shown here as the number of candidate mutants identified in each of the indicated pathways. (C) Genes required for zinc uptake identified in the screen and corresponding functions (https://www.genome.jp/dbget-bin/get_linkdb?-t+genes+gn:T00473).

Among those 80 candidates, mutations were found in genes belonging to a wide variety of functional classes (Fig. 2B), including: (i) carbohydrate transport and metabolism (*SSA_0222*, *SSA_0773*, *SSA_1261*, *SSA_1521*, *SSA_1749*, etc.), (ii) amino acid transport and metabolism (*SSA_0564*, *SSA_1043*, *SSA_1044*, *SSA_1341*-*1343*, etc.), (iii) cell division and cell envelope biogenesis (*SSA_0015*, *SSA_0655*, etc.), (iv) coenzyme metabolism (*SSA_1201*, *SSA_1202*, *SSA_1536*, etc.), (v) DNA replication (*SSA_1626*), (vi) nucleotide transport and metabolism (*SSA_0568*, *SSA_1163*, etc.), and (vii) translation, ribosomal structure and protein biosynthesis (*SSA_0820*, *SSA_1272*, *SSA_1611*, *SSA_1613*, *SSA_1895*, *SSA_1896*, *SSA_2033*, *SSA_2058*, etc.).

We also found genes involved in metal ion (zinc, iron and manganese) transport and metabolism (*SSA_0136*, *SSA_0137*, *SSA_0260*, *SSA_0261*, *SSA_1955*, *SSA_1956*, *SSA_2365*-*2367*, etc.; Table S1 and Fig. 2B,C), which are involved in many crucial biological processes and are essential for bacterial survival in the environment or in the infected host (28). In particular, zinc has recently been shown to be essential for pathogenic streptococci in their growth, morphology and virulence during infection (42, 43).

### Zinc is required for the promotion of *S. sanguinis* growth by *P. aeruginosa*

To further probe the basis for the enhanced *S. sanguinis* growth mediated by *P. aeruginosa*, we focused on the role of zinc importers identified in our screen (SSA_0136, SSA_0137, SSA_0260 and SSA_0261; Fig. 2C). In *S. sanguinis* SK36, the *SSA_0136* and *SSA_0137* genes encode components of an Adc zinc ATP-binding cassette (ABC) transporter, which is involved in zinc uptake in several pathogenic streptococci. The *SSA_0136* and *SSA_0137* genes are called *adcC* and *adcB*, respectively (42, 45). The *SSA_0260* gene, also termed SsaB (*S. sanguinis* adhesin B) (56), is an LraI family lipoprotein and serves as the substrate-binding protein for a Mn/Zn ABC import system (57). The *SSA_0261* gene, also named *ssaC*, is located upstream of *SSA_0260* and encodes a putative Mn/Zn ABC transporter permease (UniProt: A3CKL5). The *SSA_0260* and *SSA_0261* genes are predicted to be co-transcribed.

To validate our screen results, the four zinc transporter mutants were further tested in the coculture assay with *P. aeruginosa* PAO1. We found that the *SSA_0136* and *SSA _0137* mutants showed a ~2-log reduction in growth enhancement relative to that of the wild-type *S. sanguinis* SK36, while the *SSA_0260* and *SSA_0261* mutants displayed a 1-log reduction compared the wild-type *S. sanguinis* SK36 (Fig. 3A). In addition, the four mutants were also tested in monoculture to determine their growth behavior; the results revealed that these mutants grew like *S. sanguinis* WT (Fig. S2), suggesting that the growth enhancement deficiency observed for these mutants was not due to a general growth defect.

**Figure 3.**
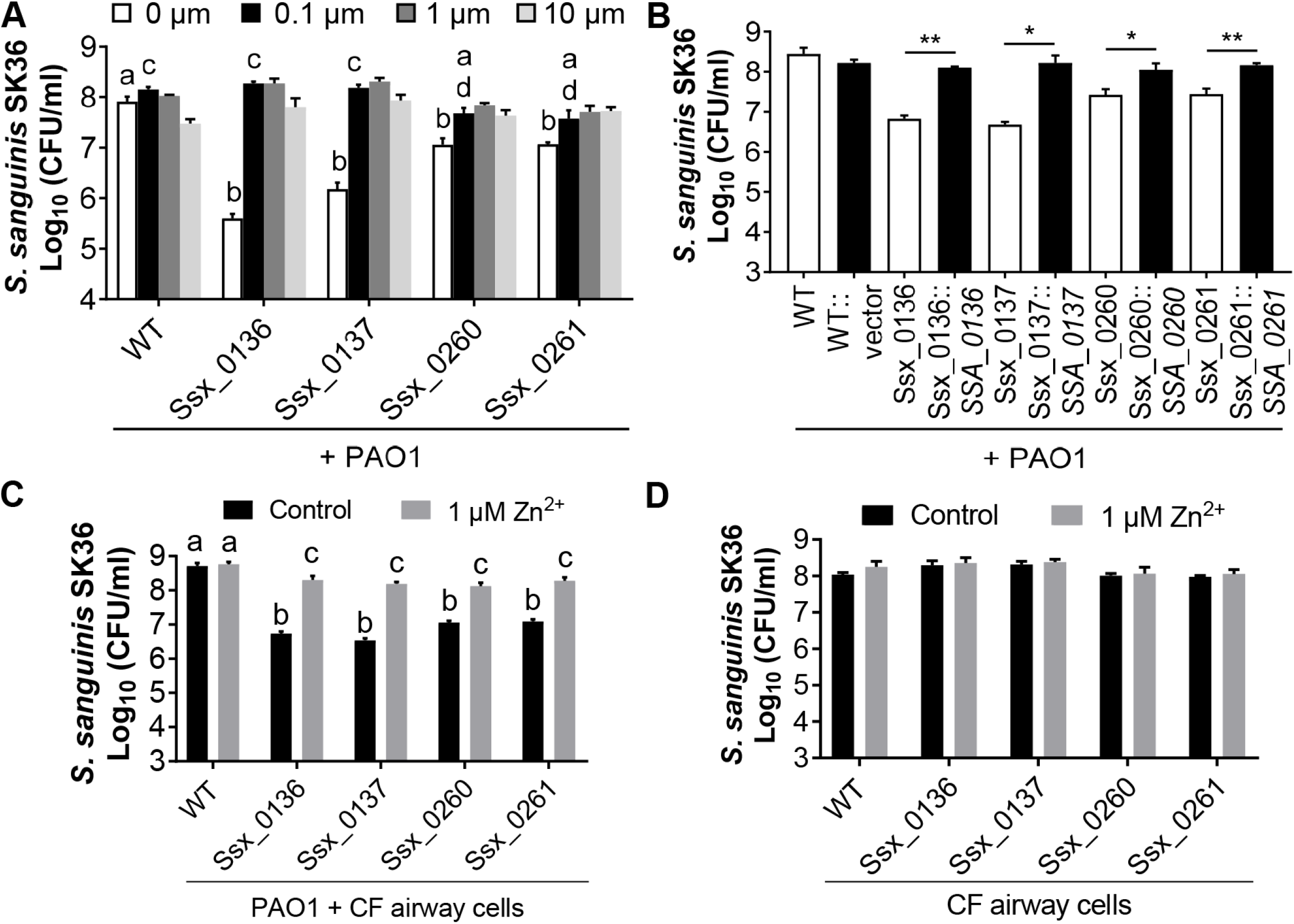
Zinc is required for the *P. aeruginosa*-induced enhancement of *S. sanguinis* growth. (A) Growth of wild-type *S. sanguinis* and zinc transporter mutants with *P. aeruginosa* (expressed as CFU/ml) in media with or without addition of zinc at the indicated concentration. Statistical significance was assessed by one-way ANOVA with a Turkey’s multiple comparison, and different letters indicates statistically significant differences (*P* < 0.05). Identical letters indicate no significant difference. In this and all subsequent panels, the *SSA* designation indicates the wild-type gene, while the Ssx designation indicates a mutation in that gene, using the convention reported in the original description of these mutant strains (55). (B) Complementation assays with the zinc transporter mutant strains. Significant differences in growth compared to the uncomplemented vector control are indicated (*, *P* < 0.05; **, *P* < 0.01, Student’s *t*-test). (C) Growth of indicated *S. sanguinis* strains with *P. aeruginosa* on CF airway cells in the presence or absence of 1 μM zinc. Statistical significance was determined by one-way ANOVA with a Turkey’s multiple comparison. Different letters indicates statistically significant differences (*P* < 0.05). Identical letters indicate no significant difference. (D) Growth of indicated *S. sanguinis* strains in monoculture on CF airway cells in the presence or absence of 1 μM zinc. Data are representative of three experiments performed in triplicate, and none of the differences are significant.

To confirm that the observed growth promotion defect was indeed caused by inactivation of the individual genes, we complemented each of the mutants with its corresponding gene under the control of an IPTG-inducible promoter, as reported (58). As shown in Fig. 3B, the complemented mutants showed growth enhancement phenotypes similar to that of wild-type *S. sanguinis* SK36, indicating that the mutation in these genes is solely responsible for the observed growth enhancement deficiency. Together, our data indicate that the functions of these genes are required for *S. sanguinis* SK36 to show enhanced growth in coculture with *P. aeruginosa*.

As zinc-ABC transporters are required for zinc acquisition (42), we reasoned that the defects in growth enhancement of these mutants during coculture might be due to the lack of zinc uptake. Consequently, the effect of supplementing additional zinc to the coculture medium was tested. We first measured the zinc levels of the medium used in our coculture conditions (MEM+L-Gln) and found the concentration of zinc is 0.024 ± 0.007 μM. Supplementation of the medium with 0.1 μM, 1 μM and 10 μM zinc chloride restored the growth enhancement phenotype of each of the mutants to wild-type levels (Fig. 3A), and growth of the wild-type *S. sanguinis* SK36 was slightly promoted upon addition of 0.1 μM and 1 μM zinc (Fig. 3A). There was no significant difference in *P. aeruginosa* growth in monoculture or coculture in the zinc-amended and unamended media (Fig. S3). These results indicate that zinc is required for efficient growth enhancement of *S. sanguinis* during coculture with *P. aeruginosa*.

Notably, zinc added to 10 μM slightly inhibited the growth of wild-type *S. sanguinis* SK36 and the SSA_0136 and SSA _0137 mutants, but not the SSA_0260 and SSA_0261 mutants (Fig. 3A), suggesting that high concentrations of zinc are toxic to *S. sanguinis*, that these zinc transporters play an important role in zinc acquisition, and may have distinct affinities for zinc.

As mentioned above, *P. aeruginosa* can also promote *S. sanguinis* SK36 growth on airway cells (Fig. 1D). To further investigate the role of zinc in the *P. aeruginosa*-induced growth enhancement, we extended our observation to cocultures on CFBE monolayers. The wild-type *S. sanguinis* SK36 strain and zinc transporter mutants were cocultured with *P. aeruginosa* with 1 μM zinc chloride added to the medium. Similar to the observations in our coculture model on plastic, the zinc transporter mutants exhibited reduced growth enhancement, and supplementing the medium with 1 μM zinc rescued the growth enhancement defect of these mutants (Fig. 3C). It is worth noting that there was no significant difference in growth of the wild-type *S. sanguinis* SK36 and the zinc transporter mutants on CFBE monolayers with or without addition of zinc (Fig. 3D). The growth of *S. sanguinis* was enhanced when cultured with CFBE cells even in the absence of zinc (Fig. 1D), indicating that CFBE monolayers produce factor(s) other than zinc that can enhance the growth of *S. sanguinis*. Taken together, these data demonstrate that enhanced *S. sanguinis* growth mediated by *P. aeruginosa* requires the zinc importers of *S. sanguinis* SK36.

### Coculture with *P. aeruginosa* upregulates zinc transporter gene expression in *S. sanguinis*

We next examined the expression patterns of zinc transporter genes in both *S. sanguinis* and *P. aeruginosa* using qRT-PCR. As shown above in Fig. 1A, in the first 2 hours of coculture with *P. aeruginosa* PAO1, *S. sanguinis* did not show a significant enhancement in growth. The mixed cultures showed a significant trend toward enhanced *S. sanguinis* growth and exhibited a robust increase in growth at 6 h. We saw no differences between the pure and the mixed cultures when we examined the growth kinetics of *P. aeruginosa* PAO1 over the course of 24 h. As a consequence, qRT-PCR studies were performed on samples of 2 h and 6 h for both monocultures and cocultures.

At 2 h, *S. sanguinis* zinc transporter genes *SSA_0136* and *SSA_0137* were upregulated 2.2- and 2.3-fold in coculture compared to *S. sanguinis* monoculture (Fig. 4A), respectively, while the expression of *SSA_0260* and *SSA_0261* genes were not significantly changed at this time point (Fig. 4A). At 6 h, *S. sanguinis* expression of all of the four zinc transporter genes was upregulated (4.4-fold for *SSA_0136*; 4.7-fold for *SSA_0137*; 8.2-fold for *SSA_0260*; 4.3-fold for *SSA_0261*) when co-cultured with *P. aeruginosa* (Fig. 4A). In contrast, the addition of exogenous zinc (1 μM) to the coculture medium reduced the expression of *S. sanguinis* zinc transporter genes to the same level as monocultures with zinc supplementation (Fig. 4B). The expression of *P. aeruginosa* PAO1 zinc uptake and regulator genes (*zur*, *znuA*, *cntI*, *cntO*) was minimally impacted by the presence of *S. sanguinis* compared to *P. aeruginosa* PAO1 alone at either 2 h or 6 h (Fig. S4), although the zinc transporters of *P. aeruginosa* PAO1 were significantly upregulated in monoculture and coculture at 6 h compared to 2 h (Fig. S4). Notably, the robust increase in zinc transporter gene expression by *S. sanguinis* at 6 h corresponds to the induction of *P. aeruginosa* uptake systems at this same time point. Taken together, our studies are consistent with a model that *S. sanguinis* zinc transporter genes are induced by zinc deficiency when *S. sanguinis* is grown in coculture with *P. aeruginosa*, presumably due to zinc competition between these organisms.

**Figure 4.**
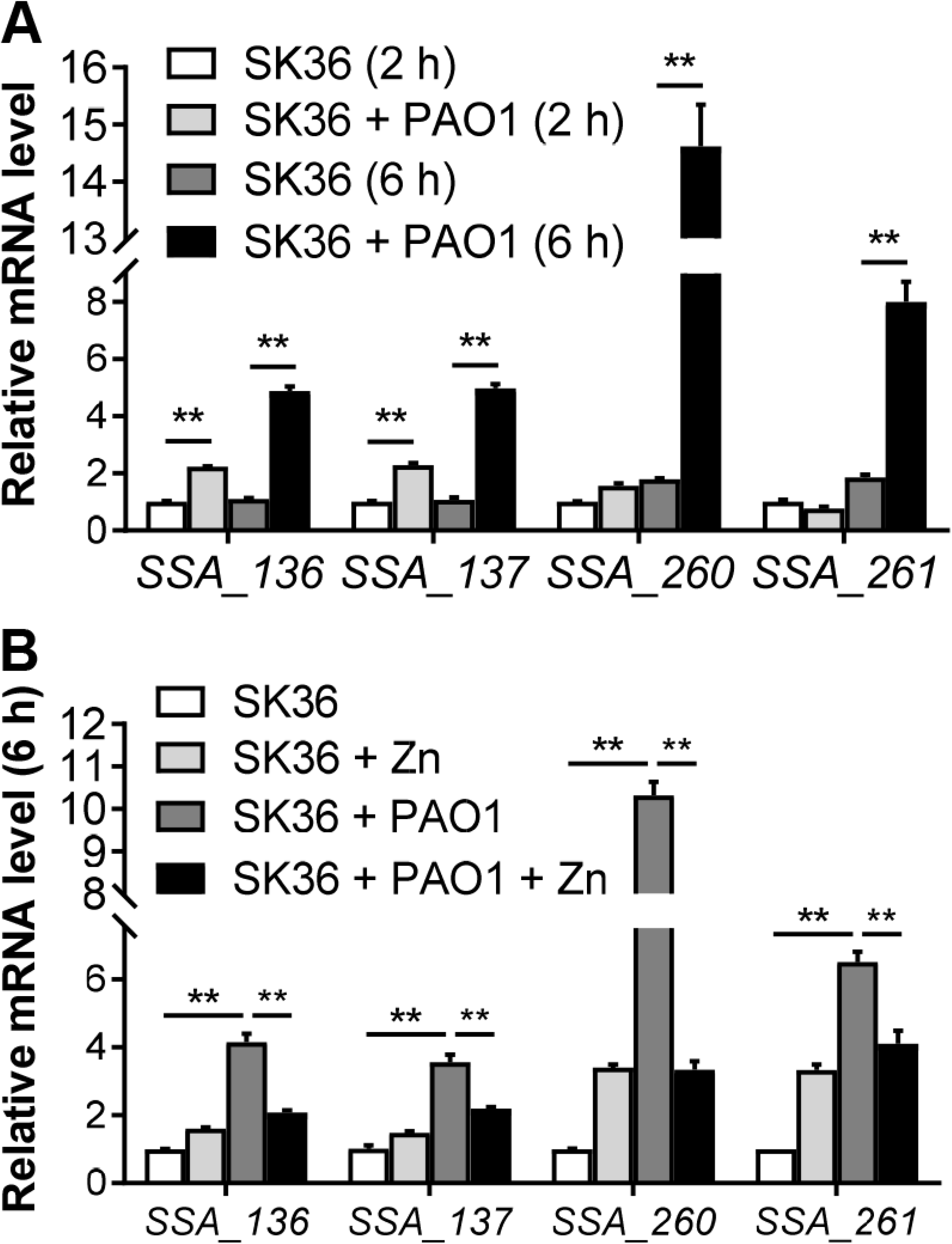
*S. sanguinis* zinc transporter genes are upregulated in the presence of *P. aeruginosa*. (A) Relative mRNA expression of *S. sanguinis* zinc transporter genes in monoculture and coculture with *P. aeruginosa* at 2 h and 6 h. The relative mRNA expression was measured using qRT-PCR, normalized to the expression of the *gyrA* control, and calculated using the 2^−ΔΔCT^ method setting the value of SK36 (2 h) as one. (B) Relative expression of *S. sanguinis* zinc transporter genes at 6 h in monoculture and coculture with *P. aeruginosa* with or without zinc (1 μM) supplementation. Error bars represent deviations of the means. ANOVA with Turkey’s multiple comparison test was used for statistical analysis (**, *P* < 0.01).

### Mutations in zinc-related genes of *P. aeruginosa* promote growth of *S. sanguinis* in coculture

Our data suggest that in coculture, *P. aeruginosa* and *Streptococcus* compete for zinc. As shown above, *S. sanguinis* SK36 mutants defective in zinc uptake show less robust viability when grown in coculture with *P. aeruginosa*. Thus, we would predict that mutations in zinc uptake in *P. aeruginosa* would result in *S. sanguinis* SK36 more effectively competing for zinc, likely reflected by an increase in viable counts of *S. sanguinis*.

We first assessed the impact of mutating zinc transport systems of *P. aeruginosa* PAO1 by testing strains with mutations in the *cntI*, *znuA* and *cntI-O* genes (Fig. 5A-C). In all cases, coculture with the *P. aeruginosa* zinc transport mutants resulted in a modest enhancement (~2-3-fold) of growth of *S. sanguinis* SK36 compared to coculture with the wild-type *P. aeruginosa* PAO1. One confounding factor in the interpretation of these results is that all of the *P. aeruginosa* zinc transport mutants show a small but consistent ~50% reduction in growth in coculture, which may be due to the lack of effective zinc transport or other anticipated defects, as reported (35, 37).

**Figure 5.**
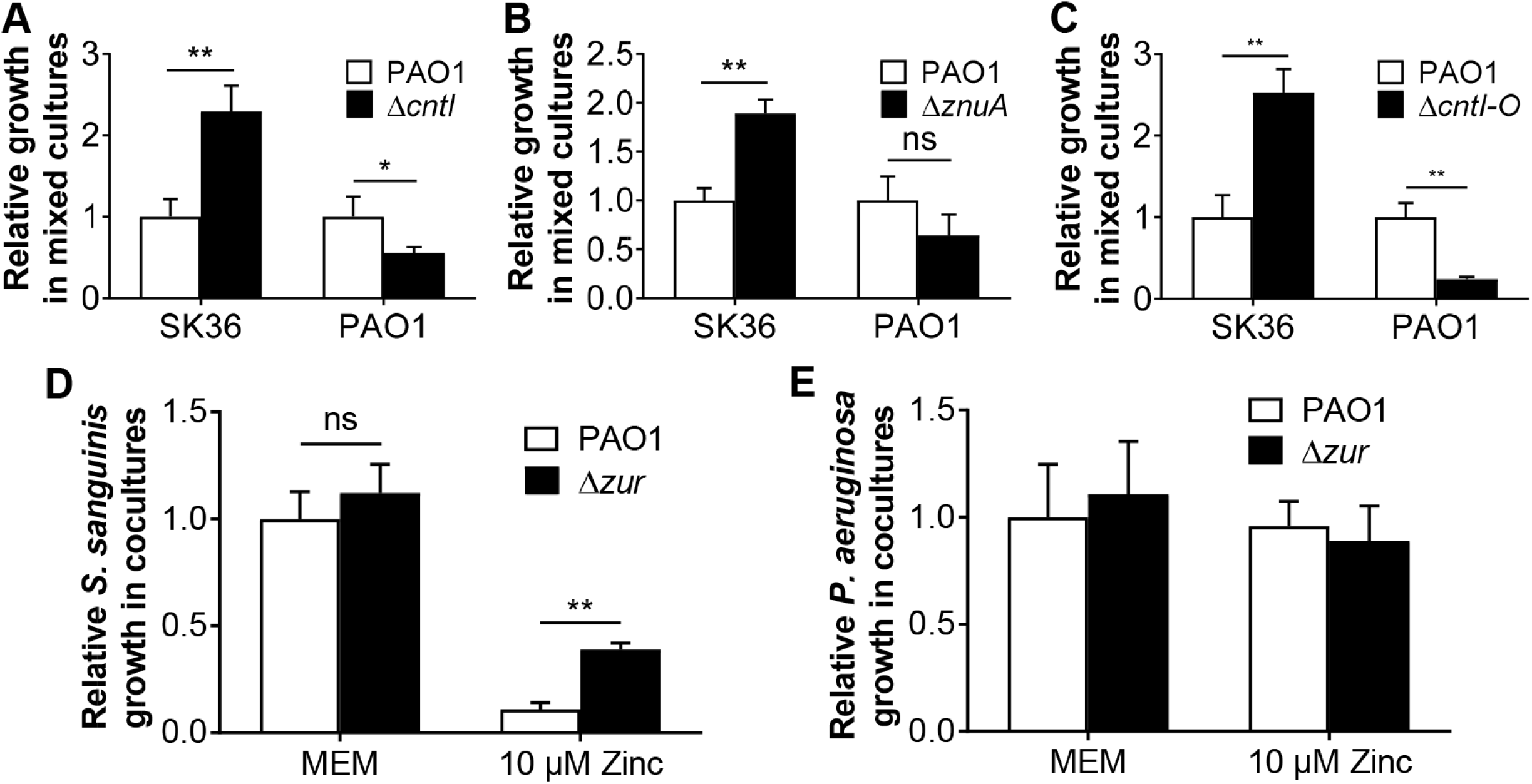
Analysis of zinc transporter mutants. (A-C) Growth of *S. sanguinis* SK36 with *P. aeruginosa* PAO1 wild type and indicated mutant strains in coculture. The strains tested carry deletion mutations in the *cntI* (A), *znuA* (B) and *cntI-O* (C) genes of *P. aeruginosa* PAO1. (D-E) Growth of *S. sanguinis* SK36 (D) and *P. aeruginosa* wild-type and ΔzurA mutants (E) in coculture assays in MEM (no added zinc) or with 10 μM of added zinc. ANOVA with Turkey’s multiple comparison test was used for statistical analysis (**, *P* < 0.01) in panel D. There were no significant differences in panel E. SK36, *S. sanguinis* SK36 and PAO1, *P. aeruginosa* PAO1.

In *P. aeruginosa*, the expression of zinc importer genes is controlled by the Zur protein, a zinc responsive repressor (36), and loss of Zur results in a higher intracellular zinc concentration in *P. aeruginosa* (37). We generated a *P. aeruginosa* PAO1 Δ*zur* mutant and cocultured this mutant with *S. sanguinis* SK36 with or without zinc addition. Coculture of the wild-type *P. aeruginosa* PAO1 with *S. sanguinis* served as the control. In the absence of added zinc, there was no difference in the growth of *S. sanguinis* SK36 in the presence of the wild-type versus *P. aeruginosa* PAO1 Δ*zur* mutant (Fig. 5D-E). As described above, in mixed cultures of *S. sanguinis* and wild-type *P. aeruginosa* PAO1, we found excessive zinc concentration (10 μM) led to a 9-fold decline in *S. sanguinis* growth compared to cells grown without zinc supplementation (Fig. 3A and Fig. 5D). In contrast, mixed cultures of *S. sanguinis* and *P. aeruginosa* PAO1 Δ*zur* mutant in the presence of 10 μM zinc resulted in a 3.5-fold increase in *S. sanguinis* growth compared to that observed for coculture with WT *P. aeruginosa* PAO1 (Fig. 5D). Given the protective action of the zinc hyper-accumulating *P. aeruginosa* PAO1 Δ*zur* mutant towards *S. sanguinis* SK36, these results further support the idea that *P. aeruginosa* and *S. sanguinis* compete for zinc when grown in coculture.

### The relationship between sputum zinc concentration and the relative abundance of *Streptococcus* and *Pseudomonas*

In previous studies we had collected sputum samples from patients with CF and assessed the relative abundance of *Streptococcus* and *Pseudomonas* in these samples (59). These same samples were analyzed by ICP-MS, as reported (59, 60), to determine the concentration of zinc in the sputum. We observed that the concentration of zinc in these samples ranged from 4.8-145 μM with a median of 36.4 μM (n = 118 sputum samples; Fig. S5).

We examined the relationship between zinc concentration and the relative abundance of *Streptococcus* and *Pseudomonas* in these samples and observed that Streptococcus dominated the sputum samples when the zinc concentration was low (Fig. 6, orange dots), but that the relative abundance of *Pseudomonas* was higher in samples with high zinc levels (Fig. 6, blue dots). Although measures like Pearson or Spearman correlation have been used to quantify this type of relationship and attach statistical significance, the compositional nature of bacterial community abundance suggests that these statistical approaches are inappropriate (61, 62). Therefore, we interpret *in vivo* associations observed between *Streptococcus* abundance*, Pseudomonas* abundance and zinc concentration as broadly consistent with the hypothesis that they are related without attaching a specific measure of statistical significance.

**Figure 6.**
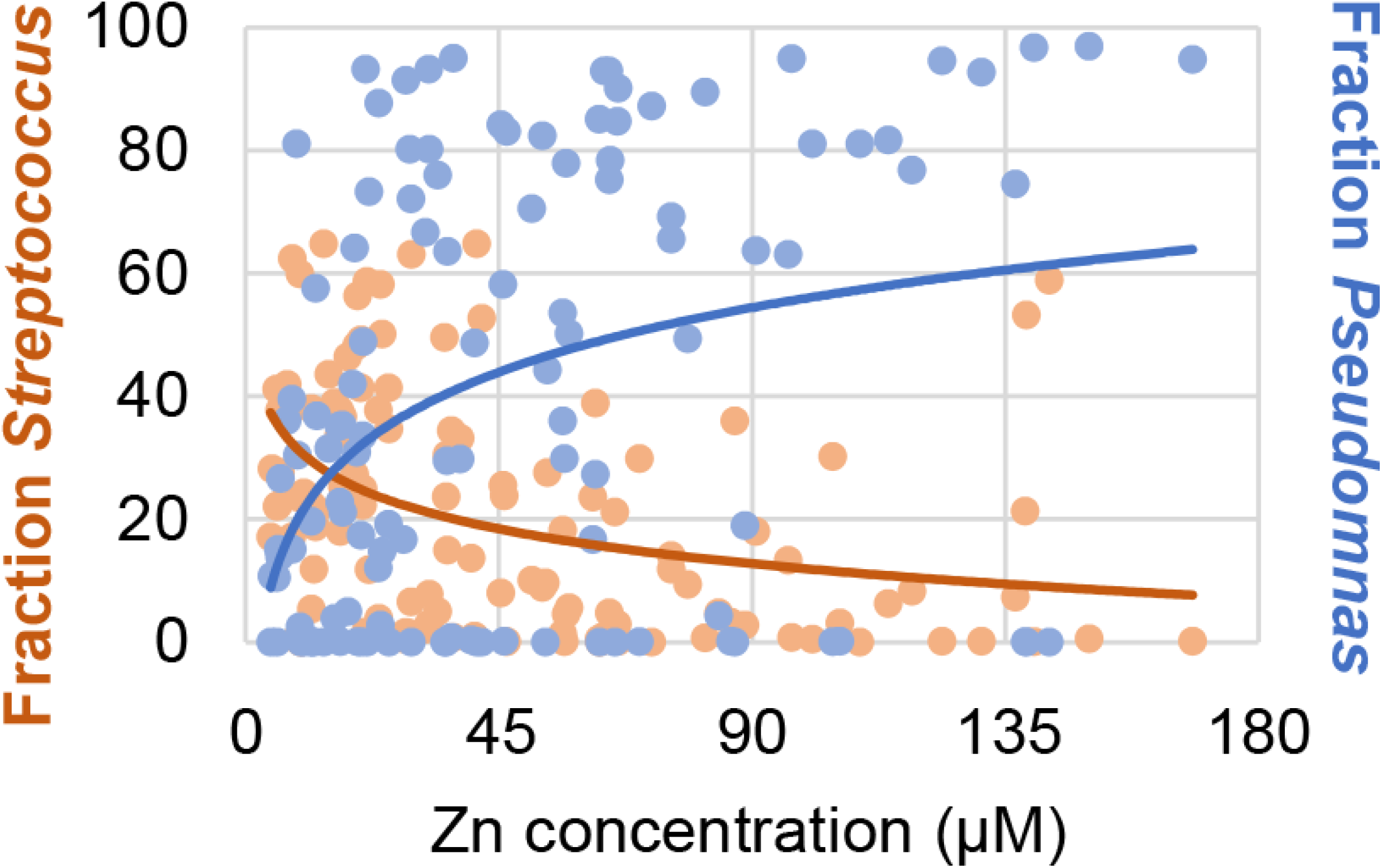
The relationship between sputum zinc concentration and the relative abundance of *Streptococcus* and *Pseudomonas*. The relative abundance of *Streptococcus* (orange dots, left Y axis) and *Pseudomonas* (blue dots, right Y axis) in each sputum sample (indicated as a fraction of 100%) is plotted versus the concentration of zinc in the corresponding sample (expressed as μM zinc on the X-axis). Total zinc was measured by ICP-MS analysis of nitrate acid-dissolved samples and normalized to the volume of the sample.

### Physiologically relevant levels of zinc impact the competition between *Streptococcus* and *Pseudomonas*

Given the observations described above, we decided to assess how varying zinc concentrations might impact the competition between various *Streptococcus* species found in the CF airway and *P. aeruginosa* (Fig. 7). For clinical isolates of *S. intermedius* (two isolates), S*. constellatus*, *S. parasanguinis* and *S. salivarius*, all of which have been found in the CF airway (11), increasing zinc across a range of concentrations measured in sputum resulted in progressively lower viability of many of these streptococci when grown in monoculture (Fig. 7A) or in coculture with *P. aerugino*sa (Fig. 7B). Thus, many of these strains appear to be sensitive to clinically-relevant concentrations of zinc.

**Figure 7.**
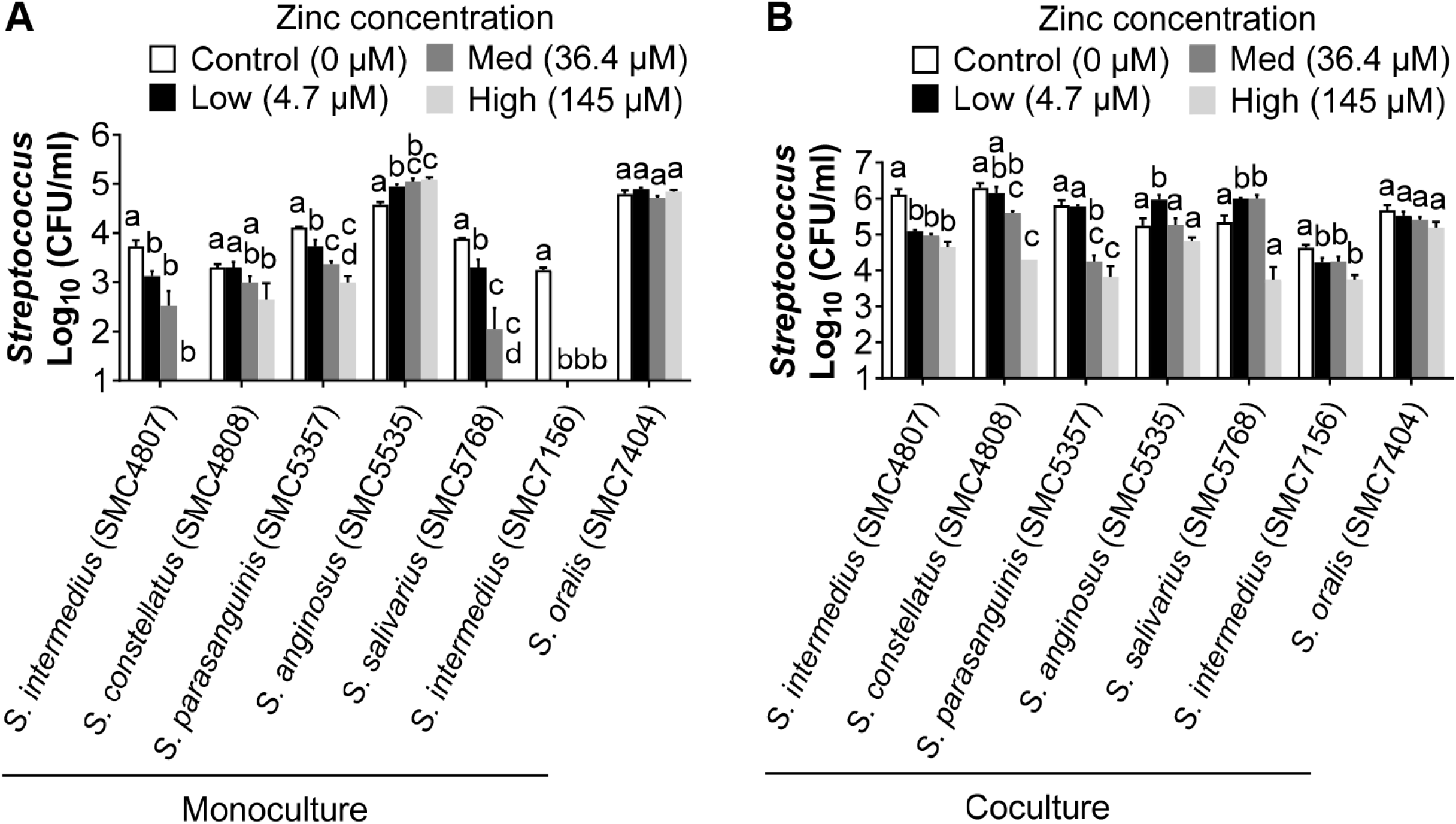
Impact of zinc level on viability of *Streptococcus* in monoculture and coculture with *Pseudomonas*. Growth of the indicated streptococci (expressed as CFU/ml) when grown in monoculture (A) or coculture with *P. aeruginosa* PAO1 (B) at the indicated concentration of supplemented zinc chloride. The 0 addition has only the zinc present in the medium (0.024 ± 0.007 μM). The other zinc additions were based on the data shown in Fig. S5 and indicate the lowest (4.7 μM), median (36.4 μM) and highest concentration (145 μM) of zinc found in the 118 sputum samples analysed. Statistical significance was determined by one-way ANOVA with a Turkey’s multiple comparison. Different letters indicates statistically significant differences (*P* < 0.05). Identical letters indicate no significant difference.

Interestingly, two isolates (*S. anginosus* and *S. oralis*) showed robust growth even at the highest level of zinc tested, indicating that these strains have increased tolerance to zinc for reasons we do not understand. Interestingly, the data in Figure 6 show that some patients with high levels of zinc in their sputum also have high relative abundance of *Streptococcus*; perhaps these patients harbor zinc-resistant streptococci.

## DISCUSSION

In this study, we characterized the interaction between *P. aeruginosa* and *S. sanguinis* SK36 in a dual species coculture model system. We demonstrated that zinc uptake by *S. sanguinis* SK36 was necessary for the *P. aeruginosa*-mediated promotion of *S. sanguinis* SK36 growth, and that *P. aeruginosa* competed with *S. sanguinis* SK36 for zinc during cocultivation. Additionally, we described a new association between zinc levels and the abundance of *Pseudomonas* and *Streptococcus* in CF sputum, thus highlighting the potential role of zinc in the interaction between these CF pathogens, as well as a potential role for this metal in shaping microbiome dynamics in the context of polymicrobial CF airway infections.

We report that co-cultivation with *P. aeruginosa* results in enhanced growth of *S. sanguinis* SK36 on either plastic or CF-derived airway cells, while *P. aeruginosa* growth is relatively unaffected during coculture. These data are consistent with previous reports from our group and others that coculture of streptococci with *P. aeruginosa* promotes the growth of streptococci, but with no obvious benefit to *P. aeruginosa* growth (8–10). Interestingly, in a study examining the interactions between *P. aeruginosa* and oral streptococci, including *S. sanguinis* (12, 63), *P. aeruginosa* growth was inhibited when streptococci were grown as a pioneer colonizer; streptococci can produce hydrogen peroxide (H_2_O_2_) to react with excess nitrite in the medium to generate reactive nitrogenous intermediates (RNI) for the inhibition of *P. aeruginosa* growth (12, 13, 63). We note that *P. aeruginosa* was inoculated in combination with *S. sanguinis* in our study under conditions that differ from these previous reports, and we did not observe streptococcus-mediated inhibition of *P. aeruginosa*. These observations suggest that the relationships between *P. aeruginosa* and streptococci are complex and likely are influenced by metabolic/environmental factors and colonization sequence. Understanding how various *in vitro* models reflect dynamic *in vivo* environments will contribute to efficiently synthesizing and better understanding the data obtained from various laboratories.

By screening a genome-wide non-essential gene mutant library of *S. sanguinis* SK36, we discovered 80 mutants with attenuation in *P. aeruginosa*-mediated growth enhancement (summarized in Table S1). The large number of genes (3.5% of the *S. sanguinis* SK36 genome) found to be involved in this interspecies interaction and the variety of functions performed by these gene products provide new insights into the mechanism(s) of interaction between these two pathogens. Among the candidate genes we identified in our screen, we focused on a set of genes involved in the import of zinc. We demonstrate that the ability to obtain zinc as one factor contributing to the *P. aeruginosa*-induced enhancement of *S. sanguinis* SK36 growth, both in the presence and absence of human airway cells. Future studies will focus on the other genes identified in our screen.

Our observations here raise the question of whether zinc is a factor that drives growth enhancement of streptococci in the presence of *P. aeruginosa*, or whether efficient uptake of zinc is required to allow the growth-promoting factors produced by *P. aeruginosa* to exert their effect. We favor the latter model, in large part because we show that adding additional zinc to monocultures of *S. sanguinis* SK36 or other streptococci has, at best, a very modest effect on growth of the streptococci. That is, it does not appear that the streptococci are obtaining zinc from *P. aeruginosa* to enhance streptococcal growth. Instead, we observed that zinc starvation might be triggered by competition for this metal between *P. aeruginosa* and *S. sanguinis* SK36. Indeed, the elevated expression of the zinc transporters of wild-type *S. sanguinis* SK36 in mixed cultures, and the lack of such induction with zinc supplementation in the coculture medium, indicates that *S. sanguinis* SK36 becomes zinc-starved during its interaction with *P. aeruginosa*. In contrast, there was no significant change in the expression of *P. aeruginosa* zinc transporters under coculture conditions, indicating that *P. aeruginosa* is not lacking for this metal under these conditions; thus it is unlikely that *S. sanguinis* is “stealing” the zinc in the culture from *P. aeruginosa*. Taken together, we argue that zinc provided by *P. aeruginosa* is not a key factor enhancing streptococcal growth in coculture.

An intriguing observation here is the relationship between the sputum zinc levels and the relative fraction of *Pseudomonas* and *Streptococcus* in the CF sputum. In the human body, the zinc concentration varies among different tissues and total zinc concentration in induced sputum from control patients is around 1 μM (50 μg/l) (42, 64), although higher levels of total sputum zinc have been reported in patients with CF (48), a finding consistent with our sputum measurements here. Furthermore, the zinc-sequestering protein calprotectin is present in CF sputum in high concentrations (65, 66), potentially resulting in limited bioavailability of zinc for CF pathogens like *P. aeruginosa* (49). Thus, at present it is difficult to conclude how much of the increased zinc measured in CF sputum is actually available to the microbes; knowing the answer to this question is key to understanding disease progression. For example, based on *in vitro* studies, efficient zinc uptake is critical for *P. aeruginosa* to express several virulence traits associated with lung colonization, including swarming, swimming motility and the ability to form biofilms (49). It has been shown that high concentrations of total zinc are correlated with airway inflammation (48), indicating that perhaps *P. aeruginosa* can access some of the large pool of zinc in some circumstances. Furthermore, sputum zinc levels were found to decrease following antibiotic treatment of CF exacerbation (48, 49), again suggestive that loss of access to zinc reduces virulence. Interestingly, we found a possible relationship between the concentrations of zinc and relative abundance of *Pseudomonas* and *Streptococcus*, however, because of the relative abundance data available for this analysis, determining a statistical correlation is difficult. Nevertheless, we do note that at higher zinc measured concentrations, the relative abundance of *Streptococcus* appears to be lower than at low concentrations of this metal, which may be related to the toxicity observed when *Streptococcus* is grown in high levels of zinc, resulting in impaired growth or death (67). These data do suggest that increased total zinc may be associated with increased bioavailable zinc. A more detailed analysis of the physiology of zinc metabolism in CF sputum and measurements of the bioavailabilty of this metal will be required to definitely address the questions raised here, and these questions should be addressed in future studies.

## MATERIALS AND METHODS

### Bacterial strains, plasmids and culture conditions

Bacterial strains and plasmids used in this study are listed in Table S2. *S. sanguinis* SK36 and other clinical streptococcal isolates were grown statically in Bacto™ Todd-Hewitt (TH) broth supplemented with 0.5% (w/v) yeast extract (THY), or on Trypticase™ soy agar plates supplemented with 5% (v/v) defibrinated sheep blood (blood agar) at 37ºC with 5% CO_2_. *P. aeruginosa* and *Escherichia coli* strains were grown in lysogeny broth (LB) medium (68) with shaking or on LB agar at 37ºC unless otherwise noted. As indicated, the following antibiotics and concentrations were used: 500 μg/ml kanamycin and 200 μg/ml spectinomycin for *S. sanguinis*; 50 μg/ml gentamycin and 150 μg/ml carbenicillin for *P. aeruginosa*; 10 μg/ml gentamicin, 50 μg/ml carbenicillin and 100 μg/ml spectinomycin for *E. coli*. For IPTG-inducible plasmids, IPTG was added to cultures to a 100 μM final concentration.

### Coculture assays

Coculture assays were performed as previously described with minor modifications (8). Briefly, overnight cultures of *P. aeruginosa* and *Streptococcus* spp. were centrifuged at 13,000 × *g* for 3 min, washed twice with phosphate buffered saline (PBS), and resuspended in minimal essential medium (MEM) supplemented with 2 mM L-glutamine (MEM+L-Gln). For coculture samples, *P. aeruginosa* inoculum was prepared to an OD_600_ of 0.05 and *Streptococcus* spp. inoculum was prepared to an OD_600_ of 0.001 in MEM+L-Gln. For monoculture controls, the inocula for *P. aeruginosa* or *Streptococcus* spp. were prepared to the same OD_600_ as for the coculture samples. Three wells of a 96-well deep well plate were inoculated per monoculture and coculture condition with 400 µl per well. Culture plates were then incubated statically at 37°C with 5% CO_2_ for 2 h, at which point the unattached planktonic cells were removed by aspiration and 400 µl of fresh MEM+L-Gln was once again added to each well. The cultures were then incubated for an additional 20 h, and both planktonic and biofilm cells were harvested together using a 96 pin replicator. Bacterial growth was determined by 10-fold serial dilutions in PBS and plated in 3 µl aliquots on *Pseudomonas* isolation agar (PIA) or blood agar supplemented with 10 µg/ml neomycin and 10 µg/ml polymixin B (SBA) for *P. aeruginosa* and *Streptococcus* spp. selective growth, respectively. After overnight incubation, bacterial colonies were counted and the colony forming units (CFU) per ml of culture were determined.

### Genetic screen of the SK36 library for mutants defective in growth enhancement

To investigate *S. sanguinis* SK36 genes involved in *P. aeruginosa*-mediated enhancement of growth, the *S. sanguinis* SK36 non-essential gene mutant library (55) was screened for growth enhancement defects as previously described for *P. aeruginosa* with some modifications (8). Briefly, a 96 pin replicator was used to transfer inocula from the frozen library to a 96-well plate containing 150 ml of THY broth per well. The plate was then incubated statically for 24 h at 37ºC with 5% CO_2_. The *P. aeruginosa* PAO1 culture was grown overnight in LB broth, adjusted to an OD_600_ of 0.05 in MEM+L-Gln as described above, and 400 µl of this adjusted inoculum suspension was added to each well of a 96-well deep well plate. The 96 pin replicator was then used to transfer 2-3 µl of the 24 h culture from the mutant library plate into the 96-well deep well plate containing *P. aeruginosa* PAO1. Unattached bacteria were removed after 2 h, and 400 µl of fresh MEM+L-Gln was added to each well and the plates were grown for an additional 20 h at 37°C with 5% CO_2_. The 96 pin replicator was again used to disrupt the biofilms into the planktonic fraction, and large petri dish plates containing either PIA or SBA media were spot inoculated with each culture and grown as described above. Candidates that showed low or undetectable growth based on differences compared to the wild-type *S. sanguinis* SK36 (which formed small lawns when inoculated onto an agar plate by the 96 pin replicator) were stored at −80ºC in 30% glycerol in a sterile 96-well plate. To confirm the phenotype, a second and third round of mutant screening was performed as described above.

### Kinetic growth assay

The relative ability of *S. sanguinis* SK36 and *P. aeruginosa* PAO1 to grow in the coculture model system was determined by kinetic growth assays. Briefly, *S. sanguinis* SK36 was grown in coculture with *P. aeruginosa* PAO1 as described above in a 96-well deep well plate. The cultures were grown at 37°C with 5% CO_2_ for 24 h, and were assessed for viable cell counts (CFU/ml) at seven time points: 0, 2, 4, 6, 8, 10, and 24 h. The 0 h time point corresponds to the initial inoculum. Cells were collected at the 2 h time point prior to the 2 h medium replacement. At each time point, a combination of the planktonic and biofilm cells from triplicate wells were serially diluted and plated on PIA and SBA agar media, and CFU counts determined after overnight incubation as described above.

### *P. aeruginosa* supernatant assays

To prepare *P. aeruginosa* conditioned medium, bacteria were grown in 0.5 ml per well of MEM+L-Gln in 24-well plates with media changes as described above. At the 22 h time point, the planktonic fractions were collected, centrifuged and supernatants were filter-sterilized through a 0.22 μm syringe filter. The effect of the sterile *P. aeruginosa* supernatants on the growth of *S. sanguinis* was tested on both plastic and CFBE monolayers. For experiments on plastic, supernatant with different levels of dilution in fresh MEM+L-Gln was added to *S. sanguinis* monocultures when the medium was replaced at the 2 h time point. For assays on CF airway cells, a 1/2 dilution of *P. aeruginosa* supernatant in MEM+L-Gln was supplemented with 0.4% L-arginine and gently added to each well of CF airway cells that had been cocultured with *S. sanguinis* for 1 h or 5.5 h when the medium was exchanged. *S. sanguinis* growth was evaluated after 22 h of incubation at 37°C with 5% CO_2_ as described above.

### Tissue culture cells and coculture on CFBE monolayers

The cystic fibrosis bronchial epithelial (CFBE) monolayers used in the coculture model (54, 69) are immortalized cells that overexpress ΔF508-cystic fibrosis transmembrane conductance regulator (10, 70). CFBE monolayers were grown as previously described (10, 69). In brief, the CFBE monolayers were seeded at a concentration of 100,000 cells/well in a 24-well tissue culture plate and fed every other day with MEM supplemented with 10% fetal bovine serum, 2 mM L-glutamine, 50 U/ml penicillin, 50 μg/ml streptomycin, 2 μg/ml puromycin and 5 μg/ml Plasmocin. Cells were grown at 37°C with 5% CO_2_ for 5-7 days to form a confluent monolayer and tight junctions before inoculation with bacteria. For coculture assays with mono- and dual-bacterial species, liquid cultures of *P. aeruginosa* and *Streptococcus* spp. were prepared as described above and 500 μl of bacterial inocula were gently added to triplicate wells of CFBE monolayers that had been washed twice with MEM. The cocultures were incubated at 37°C with 5% CO_2_ for 1 h, at which point unattached bacteria were removed by aspiration, 500 μl of MEM+L-Gln+0.4% L-arginine was added to each well and incubated for an additional 4.5 h. At this point, planktonic cells were removed by aspiration, and 500 µl of fresh MEM+L-Gln+0.4% L-arginine was once again added into each well. The established coculture was incubated for an additional 16.5 h. At 22 h post-inoculation, both planktonic and biofilm-grown bacteria were collected together by scraping with a pipette tip, and bacteria were serially diluted and plated on PIA and SBA plates, as described above, to identify *P. aeruginosa* and *Streptococcus* spp., respectively. Following overnight incubation, the resulting colonies were counted and the CFU/ml of the culture was determined.

### Zinc supplementation

For zinc supplementation assays, wild-type *Streptococcus* SK36 and individual mutants with growth enhancement defects, wild-type *P. aeruginosa* PAO1 and *P. aeruginosa* PAO1 zinc homeostasis associated mutants were grown as described above for monoculture and coculture assays. After 2 h, unattached bacteria were removed by aspiration, MEM+L-Gln with or without additional zinc (at the indicated concentrations diluted from 1 mM zinc chloride stock solution in MEM+L-Gln) was added to each well and cocultures were then treated as described above. Nutritional complementation on CF airway cells were performed as described for biofilms on plastic, except that MEM+L-Gln supplemented with 0.4% arginine (to enhance biofilm formation) with or without additional zinc chloride at the indicated concentration was used for medium exchange when indicated.

### Construction of mutants and complementation

In-frame deletions of *P. aeruginosa* genes were constructed by allelic exchange employing the sucrose counter-selection system with the gene replacement vector pEX18Ap (71). Mutant strains were confirmed by PCR analysis of genomic DNA. *S. sanguinis* mutants were derived from a defined mutant library described previously (55). For complementation of each targeted *S. sanguinis* gene, a suicide vector pJFP126 was used to allow for the insertion of complementing genes into an ectopic chromosomal site (*SSA_0169*) via homologous recombination and expression of each gene is under the control of an IPTG inducible promoter *hyper-spank* (8, 58). Transformation was performed essentially as described previously (72) with the competence stimulating peptide (CSP sequence: DLRGVPNPWGWIFGR) custom-synthesized by GenScript Inc. (Piscataway, NJ). Primers used for PCR amplification of selected genes are listed in Table S3.

### Expression studies

For qRT-PCR studies, the overnight culture used as inoculum was prepared as described above from which three replicates cocultures of *P. aeruginosa* and *S. sanguinis*, or the *S. sanguinis* monoculture, were prepared in 100 ml of warm MEM+L-Gln in a 250 ml flask. The inoculum of *P. aeruginosa* and *S. sanguinis* was prepared and brought to an OD_600_ of 0.05 and 0.02, respectively, as described above. Cultures were incubated at 37ºC with 5% CO_2_. After 2 h and 6 h of incubation, samples were pelleted, and bacterial cells were mechanically lysed with 10 cycles of 30 s bead beating, 30 s on ice with a 1:1 mixture of 0.1 mm and 0.5 mm glass beads. Total RNA was isolated using TRIzol and the Direct-zol™ RNA MiniPrep Kit with two times of in-column DNase I treatment according to the manufacturer’s instructions (Zymo Research, R2053). RNA purity and concentration were determined using a NanoDrop 2000 spectrophotometer (Thermo Scientific). For each sample, 1 μg of RNA was converted to cDNA using the SuperScript™ III First-Strand Synthesis System for RT-PCR (Invitrogen) and then diluted 1:50. qRT-PCR were carried out in triplicate in a StepOnePlus Real-Time PCR System (Applied Biosystems) using iTaq Universal SYBR Green Supermix (Bio-Rad). The qRT-PCR primers are listed in Table S3. Relative gene expression was calculated using the 2^−ΔΔCT^ method with DNA gyrase subunit gene *gyrA* as a normalization control for *S. sanguinis* (73, 74) and *PA2875* as the reference gene for *P. aeruginosa* (75).

### Measurement of zinc in sputum samples

Sputum samples for zinc analysis were stored at −80°C until processed. Sputum zinc was quantified by inductively coupled plasma-mass spectrometry (ICP-MS) following nitric acid digestion of organic material according to the method of Heck et al. and is expressed as micromolar zinc (60).

### Statistical Analysis

Statistical analysis was performed using GraphPad Prism 7 program and results were expressed as the mean values ± standard deviations. Unless otherwise noted, one-way analysis of variance (ANOVA) followed by Tukey’s multiple comparison test or Student’s *t*-test analysis was performed to determine statistical significance of the data. *, *P* < 0.05; **, *P* < 0.01. See figure legends or text for other specific statistical tests used.

## Acknowledgements

This work was supported by grants from the Cystic Fibrosis Foundation (OTOOLE16GO) and NIH (R37 AI83256-06) to G.A.O, a China Scholarship Council (CSC) grant (201708330005) to K.L. T.H. is supported by the DartCF CF-BBC (P30-DK117469). We acknowledge support from the CF-Research Development Program [STANTO19R0] for the CFBE cells. The ICP-MS studies were performed by the Dartmouth Trace Element Analysis Core, funded by grant P42ES007373 from the National Institute of Health. We thank Brian Jackson for performing these studies. We also thank P. Xu for sharing the *S. sanguinis* mutant library.

