## Supplementary material for "Availability of Zinc Impacts Interactions Between *Streptococcus sanguinis* and *Pseudomonas aeruginosa* in Co-culture": Table S1

| Table S1. Candidate Mutants. | | | | | | |
|---|---|---|---|---|---|---|
| Well Location | Gene ID | CFU from Initial Screen | CFU confirm-1 | CFU confirm-2 | Locus tag | Predicted Function |
| **Plate 1** | | | | | | |
| A03 | SSA_2390 | 30-300 | 30-300 | 30-300 | SSA_2390 | hypothetical protein |
| B02 | SSA_0015 | 300-1000 | 1000-3000 | 300-1000 | SSA_0015 | membrane ATPase FtsH, degrades sigma32 (integral membrane cell-division Zn metallo-peptidase) |
| C02 | SSA_0028 | 300-1000 | 3000+ | 30-300 | SSA_0028 | phosphoribosylaminoimidazole-succinocarboxamide synthase |
| C08 | SSA_0035 | 30-300 | 3000+ | 300-1000 | SSA_0035 | bifunctional phosphoribosylaminoimidazolecarboxamide formyltransferase/IMP cyclohydrolase |
| D08 | SSA_0048 | 30-300 | 3000+ | 30-300 | SSA_0048 | TetR/AcrR family transcriptional regulator |
| **Plate 2** | | | | | | |
| C12 | SSA_0136 | 30-300 | 1000-3000 | 3000+ | SSA_0136 | ABC transporter, Zn porter |
| D01 | SSA_0137 | 300-1000 | 300-1000 | 30-300 | SSA_0137 | ABC transporter (permease), Zn porter |
| **Plate 3** | | | | | | |
| C05 | SSA_0222 | 1-30 | 30-300 | 1-30 | SSA_0222 | PTS system, mannose-specific IID component |
| F05 | SSA_0260 | 1-30 | 30-300 | 1-30 | SSA_0260 | manganese/Zinc ABC transporter substrate-binding protein |
| F06 | SSA_0261 | 300-1000 | 1000-3000 | 1-30 | SSA_0261 | ABC-type Mn2+/Zn2+ transport systems, permease component |
| **Plate 4** | | | | | | |
| E04 | SSA_0342 | 1000-3000 | 1000-3000 | | SSA_0342 | pyruvate formate-lyase |
| E11 | SSA_0351 | 0 | 0 | 0 | SSA_0351 | Signal peptidase I |
| **Plate 5** | | | | | | |
| F10 | SSA_0460 | 0 | 0 | 1000-3000 | SSA_0460 | multiple antibiotic resistance operon transcription repressor (MarR) |
| **Plate 6** | | | | | | |
| G05 | SSA_0564 | 0 | 1-30 | | SSA_0564 | aminotransferase AlaT |
| G09 | SSA_0568 | 30-300 | 30-300 | 30-300 | SSA_0568 | aspartyl-tRNA synthetase 1 |
| **Plate 7** | | | | | | |
| F11 | SSA_0655 | 30-300 | 1-30 | 1-30 | SSA_0655 | cell division protein FtsA |
| **Plate 8** | | | | | | |
| G05 | SSA_0759 | 3000+ | | 1000-3000 | SSA_0759 | acetylglutamate kinase |
| H06 | SSA_0773 | 1000-3000 | 1000-3000 | 1000-3000 | SSA_0773 | PTS enzyme I |
| **Plate 9** | | | | | | |
| C03 | SSA_0802 | 3000+ | 300-1000 | 1000-3000 | SSA_0802 | hypothetical protein |
| D09 | SSA_0820 | 30-300 | 30-300 | 1-30 | SSA_0820 | ribosomal protein S21 |
| G09 | SSA_0861 | 3000+ | 3000+ | 3000+ | SSA_0861 | hypothetical protein |
| H02 | SSA_0866 | 0 | 0 | 0 | SSA_0866 | cation transporter E1-E2 family ATPase |
| **Plate 11** | | | | | | |
| E07 | SSA_1030 | 3000+ | 1000-3000 | | SSA_1030 | TetR family transcriptional regulator |
| F08 | SSA_1043 | 3000+ | 300-1000 | 30-300 | SSA_1043 | homoserine dehydrogenase |
| F09 | SSA_1044 | 3000+ | 30-300 | 30-300 | SSA_1044 | homoserine kinase |
| **Plate 12** | | | | | | |
| A03 | SSA_1072 | 3000+ | 1000-3000 | 30-300 | SSA_1072 | gamma-glutamyl kinase |
| A05 | SSA_1074 | 3000+ | 300-1000 | 1-30 | SSA_1074 | pyrroline-5-carboxylate reductase |
| H03 | SSA_1163 | 30-300 | 1-30 | 1-30 | SSA_1163 | bifunctional GMP synthase/glutamine amidotransferase protein |
| **Plate 13** | | | | | | |
| C07 | SSA_1201 | 30-300 | 30-300 | 300-1000 | SSA_1201 | phosphopantothenate--cysteine ligase |
| C08 | SSA_1202 | 30-300 | 30-300 | 300-1000 | SSA_1202 | phosphopantothenoylcysteine decarboxylase |
| C09 | SSA_1203 | 1000-3000 | 300-1000 | 1000-3000 | SSA_1203 | hypothetical protein |
| C10 | SSA_1204 | 0 | 0 | 0 | SSA_1204 | phosphoglucomutase |
| H06 | SSA_1261 | 30-300 | 3000+ | 1-30 | SSA_1261 | ribose-5-phosphate isomerase A |
| **Plate 14** | | | | | | |
| A06 | SSA_1270 | 1000-3000 | 300-1000 | 300-1000 | SSA_1270 | flavodoxin |
| A08 | SSA_1272 | 3000+ | | 1000-3000 | SSA_1272 | 50S ribosomal protein L31 type B |
| G02 | SSA_1341 | 3000+ | 3000+ | | SSA_1341 | carbamoyl phosphate synthase large subunit |
| G03 | SSA_1342 | 3000+ | 3000+ | 3000+ | SSA_1342 | carbamoyl phosphate synthase small subunit |
| G04 | SSA_1343 | 3000+ | 3000+ | 300-1000 | SSA_1343 | aspartate carbamoyltransferase catalytic subunit |
| **Plate 15** | | | | | | |
| A03 | SSA_1363 | 3000+ | 3000+ | 300-1000 | SSA_1363 | FmtA-like protein |
| B11 | SSA_1383 | 1000-3000 | 30-300 | 300-1000 | SSA_1383 | aspartate aminotransferase |
| E01 | SSA_1410 | 0 | 0 | 0 | SSA_1410 | dTDP-4-keto-6-deoxyglucose-3,5-epimerase |
| E02 | SSA_1411 | 0 | 0 | 0 | SSA_1411 | glucose-1-phosphate thymidylyltransferase |
| **Plate 16** | | | | | | |
| E03 | SSA_1507 | 0 | 0 | 0 | SSA_1507 | ABC-type lipopolysaccharide transport system, ATPase component |
| E04 | SSA_1508 | 0 | 0 | 0 | SSA_1508 | ABC-type lipopolysaccharide transport system, permease component |
| E05 | SSA_1509 | 3000+ | 1-30 | 1-30 | SSA_1509 | polysaccharide biosynthesis protein/ rhamnosyltransferase |
| E06 | SSA_1510 | 0 | 1-30 | 0 | SSA_1510 | rhamnosyltransferase |
| E07 | SSA_1511 | 0 | 0 | 0 | SSA_1511 | glycosyltransferase |
| E09 | SSA_1513 | 0 | 0 | 0 | SSA_1513 | glycosyltransferase |
| F05 | SSA_1521 | 0 | 0 | 0 | SSA_1521 | phosphoenolpyruvate carboxylase |
| G06 | SSA_1536 | 30-300 | 30-300 | 30-300 | SSA_1536 | biotin synthase |
| H04 | SSA_1547 | 300-1000 | 3000+ | 1000-3000 | SSA_1547 | HPr kinase/phosphorylase |

| | | | | | | |
|---|---|---|---|---|---|---|
| **Plate 17** | | | | | | |
| B10 | SSA_1574 | 30-300 | 30-300 | 30-300 | SSA_1574 | glycosyl transferase family protein |
| B11 | SSA_1575 | 30-300 | 30-300 | 30-300 | SSA_1575 | glycosyl transferase |
| E09 | SSA_1611 | 3000+ | | 3000+ | SSA_1611 | GTP-binding protein Era |
| E11 | SSA_1613 | 3000+ | 30-300 | 3000+ | SSA_1613 | putative metalloprotease |
| F12 | SSA_1626 | 3000+ | 300-1000 | 3000+ | SSA_1626 | DNA translocase ftsK |
| G12 | SSA_1639 | 3000+ | 30-300 | 1000-3000 | SSA_1639 | 5'-methylthioadenosine/S-adenosylhomocysteine nucleosidase |
| **Plate 19** | | | | | | |
| A05 | SSA_1749 | 30-300 | 1-30 | 30-300 | SSA_1749 | pyruvate formate-lyase-activating enzyme |
| C06 | SSA_1773 | 0 | 0 | 0 | SSA_1773 | hypothetical protein |
| **Plate 20** | | | | | | |
| A08 | SSA_1845 | 30-300 | 300-1000 | 30-300 | SSA_1845 | Serine/threonine protein kinase |
| B11 | SSA_1860 | 3000+ | 1000-3000 | | SSA_1860 | penicillin-binding protein 1A |
| E06 | SSA_1895 | 1000-3000 | | 3000+ | SSA_1895 | ribosome-binding factor A |
| E07 | SSA_1896 | 300-1000 | | 3000+ | SSA_1896 | translation initiation factor IF-2 |
| **Plate 21** | | | | | | |
| A07 | SSA_1942 | 3000+ | 30-300 | 30-300 | SSA_1942 | enoyl-CoA hydratase |
| A08 | SSA_1943 | 1-30 | 30-300 | 30-300 | SSA_1943 | aspartate kinase |
| B06 | SSA_1953 | 3 | 30-300 | 1-30 | SSA_1953 | NifU family protein |
| B07 | SSA_1954 | 1-30 | 30-300 | 1-30 | SSA_1954 | aminotransferase, class-V |
| B08 | SSA_1955 | 30-300 | 30-300 | 30-300 | SSA_1955 | ABC-type Fe-S cluster assembly transporter, permease component |
| B09 | SSA_1956 | 1-30 | 30-300 | 30-300 | SSA_1956 | ABC-type Fe-S cluster assembly transporter, ATPase component |
| D07 | SSA_1981 | 1-30 | | 3000+ | SSA_1981 | hypothetical protein |
| **Plate 22** | | | | | | |
| A04 | SSA_2033 | 3000+ | 1000-3000 | 1000-3000 | SSA_2033 | 30S ribosomal protein S9 |
| C03 | SSA_2058 | 3000+ | 300-1000 | 1000-3000 | SSA_2058 | 30S ribosomal protein S15 |
| G11 | SSA_2118 | w growth, 300 | 30-300 | 30-300 | SSA_2118 | thiamine pyrophosphokinase |
| **Plate 23** | | | | | | |
| D05 | SSA_2168 | 300-1000 | 1-30 | 0 | SSA_2168 | NAD(P)H-dependent glycerol-3-phosphate dehydrogenase |
| E08 | SSA_2185 | 0 | 0 | 0 | SSA_2185 | adenylosuccinate synthetase |
| **Plate 25** | | | | | | |
| D07 | SSA_2364 | 0 | 0 | 0 | SSA_2364 | immunodominant staphylococcal antigen A precursor |
| D08 | SSA_2365 | 0 | 0 | 0 | SSA_2365 | cobalt transport protein cbiQ |
| D09 | SSA_2366 | 0 | 0 | 0 | SSA_2366 | cobalt transporter ATP-binding subunit |
| D10 | SSA_2367 | 0 | 0 | 0 | SSA_2367 | cobalt transporter ATP-binding subunit |
| E05 | SSA_2374 | 3000+ | 1-30 | 1-30 | SSA_2374 | inositol-5-monophosphate dehydrogenase |
| **Zn/Mn-related functions** | | | | | | |
