## Supplementary material for "Availability of Zinc Impacts Interactions Between *Streptococcus sanguinis* and *Pseudomonas aeruginosa* in Co-culture": Table S2

**Table S2. Strains and plasmids used in this study.**

| Strain or plasmid | Relevant genotype or description | Source or reference |
|---|---|---|
| *Streptococcus* strains | | |
| *S. sanguinis* SK36 | WT isolate from human dental plaque; virulent in rat and rabbit models of infective endocarditis | (1) |
| Ssx_0136 | *S. sanguinis* SK36 Δ*SSA_0136*::*aphA-3*; zinc ABC transporter | (2) |
| Ssx_0137 | *S. sanguinis* SK36 Δ*SSA_0137*::*aphA-3*; $Km^r$; zinc ABC transporter permease | (2) |
| Ssx_0260 | *S. sanguinis* SK36 Δ*SSA_0260*::*aphA-3*; $Km^r$; Mn/Zn ABC transporter substrate-binding protein | (2) |
| Ssx_0261 | *S. sanguinis* SK36 Δ*SSA_0261*::*aphA-3*; $Km^r$; Mn/Zn ABC transporter permease | (2) |
| Ssx_0136::*SSA_0136* | *S. sanguinis* SK36 Δ*SSA_0136*::*aphA-3* SSA_0169::*aad9* Phyper-spank *lacZo* *SSA_0136 lacI*; $Kan^r$ $Spc^r$ | This study |
| Ssx_0137::*SSA_0137* | *S. sanguinis* SK36 Δ*SSA_0137*::*aphA*-3 SSA_0169::*aad9* Phyper-spank *lacZo* *SSA_0137 lacI*; $Kan^r$ $Spc^r$ | This study |
| Ssx_0260::*SSA_0260* | *S. sanguinis* SK36 Δ*SSA_0260*::*aphA*-3 SSA_0169::*aad9* Phyper-spank *lacZo* *SSA_0260 lacI*; $Kan^r$ $Spc^r$ | This study |
| Ssx_0261::*SSA_0261* | *S. sanguinis* SK36 Δ*SSA_0261*::*aphA*-3 SSA_0169::*aad9* Phyper-spank *lacZo* *SSA_0261 lacI*; $Kan^r$ $Spc^r$ | This study |
| SMC4807 | *S. intermedius* clinical isolate | This study |
| SMC4808 | *S. constellatus* clinical isolate | This study |
| SMC5357 | *S. parasanguinis* ATCC# 15912 | ATCC |
| SMC5535 | *S. anginosus* clinical isolate | This study |
| SMC5768 | *S. salivarius* JIM8780 | (3) |
| SMC7156 | *S. intermedius* | This study |

**Table S1 continued.**

| Strain or plasmid | Relevant genotype or description | Source or reference |
|---|---|---|
| *P. aeruginosa* strains | | |
| PAO1 | wild type | (4) |
| Δ*zur* | PAO1 with a *zur* deletion | This study |
| Δ*znuA* | PAO1 with a *znuA* deletion | This study |
| Δ*cntO*-c*ntI* | PAO1 with a *cntO*-c*ntI* deletion | This study |
| *Escherichia coli* strains | | |
| S17-1 (λpir) | *thi pro hdsR hdsM* $^{+}$ *recA*; chromosomal insertion of RP4-2 (Tc::Mu Km::Tn*7*) | (5) |
| **Plasmids** | | |
| pEX18Ap | Gene replacement vector; Amp$^{r}$, *oriT*$^{+}$, *sacB*$^{+}$ | (6) |
| pJFP126 | Spc$^{r}$; derivative of pJFP106 containing Phyper-spank l*acZo lacI* from pDR111 | (7) |
