## Supplementary material for "Availability of Zinc Impacts Interactions Between *Streptococcus sanguinis* and *Pseudomonas aeruginosa* in Co-culture": Table S3

**Table S3. Primers used in this study.**

| Primer | Sequence (5' to 3') | Use |
|---|---|---|
| SSA_0136-com-F | CCCAAGCTTTGGAAGCCTTAGTGGGAGAG | Complementing Ssx_0136 mutant |
| SSA_0136-com-R | ACATGCATGCGGCAATAACTGCCAAGAAGG | |
| SSA_0137-com-F | CCCAAGCTTCTTGGCGCTGTTTCAATGTG | Complementing Ssx_0137 mutant |
| SSA_0137-com-R | ACATGCATGCTAGACCTGCTAATAGTAAGC | |
| SSA_0260-com-F | ACATGCATGCTTTGATGGAGGACTCCCATG | Complementing Ssx_0260 mutant |
| SSA_0260-com-R | ACATGCATGCCCATGTGACGCTCCTTCTCC | |
| SSA_0261-com-F | CCCAAGCTTCTGATTGCTTTCGGTCCTAC | Complementing Ssx_0261 mutant |
| SSA_0261-com-R | CCCAAGCTTGACCGGAGCAGGCAAAGAG | |
| zur-dou-F | AACTTGCGCAGGTGCATGCC | Confirming Δ*zur* deletion mutant |
| zur-dou-R | TTGAAGGACACGCCGACCTG | |
| znuA-dou-F | ACCGCGAGGATATCGTAGGC | Confirming Δ*znuA* deletion mutant |
| znuA-dou-R | TCAGGCAGCAACAGACAACC | |
| cntO-dou-F | GCCAGGAAATCATGTACGCC | Confirming Δ*cntO*-*cntI* deletion mutant |
| cntI-dou-R | GCCGAAGCCTATGATCCCTG | |
| gyrA-RT-F | GAAGGATGAAGACGAGTT | Real time PCR |
| gyrA-RT-R | ATTGAAGCGGACAGAATA | |
| SSA_0136-RT-F | GGCTGTCTTGATGATTACG | Real time PCR |
| SSA_0136-RT-F | TTACGGACCAGATGAATGT | |
| SSA_0137-RT-F | GAAATCGGGACGGCTATTCTTA | Real time PCR |
| SSA_0137-RT-R | GAGCTAGAACTCTTGCCCTTAC | |
| SSA_0260-RT-F | AATGATTGTGACCAGTGA | Real time PCR |
| SSA_0260-RT-R | TTCTTCTTCGGTGTTGAT | |
| SSA_0261-RT-F | TCTTAATCATCATCCTCTT | Real time PCR |

**Table S2., continued**

| Primer | Sequence (5' to 3') | Use |
|---|---|---|
| SSA_0261-RT-R | AGAATCATCAGCAGATAG | Real time PCR |
| PA2875-RT-F | AGTTTCCAGCGCATCCAGTT | Real time PCR |
| PA2875-RT-R | CGGGATGGAAGACGAATTG | |
| cntO-RT-F | CGAGCGTTCCTATAACAA | Real time PCR |
| cntO-RT-F | TCAGTAGTTCAGGGTCAG | |
| cntI-RT-F | TGTACCTGGTCTGCTATT | Real time PCR |
| cntI-RT-R | AAGAGGATGACGAAGAAC | |
| znuA-RT-F | GTGGACGCTAACGGCTAT | Real time PCR |
| znuA-RT-R | TCAGAGCTTTTCCAGACAG | |
| zur-RT-F | TTCGTCGGCTGCAACAA | Real time PCR |
| zur-RT-R | GGCTGATATCCGGCTGTTC | |
