## Supplemental Figures for "Availability of Zinc Impacts Interactions Between *Streptococcus sanguinis* and *Pseudomonas aeruginosa* in Co-culture"

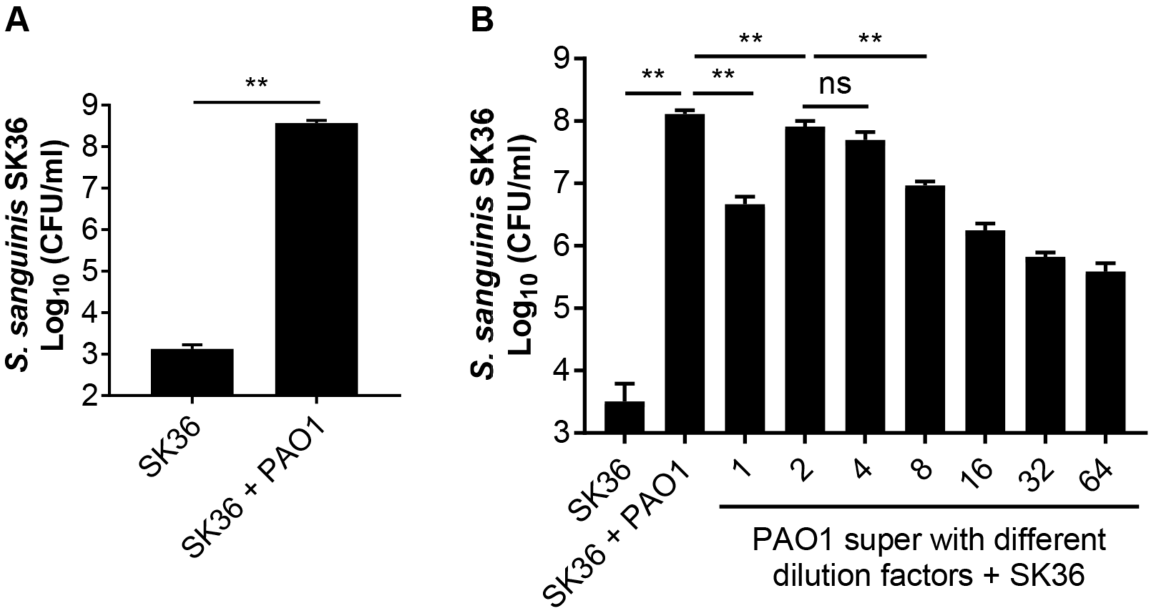


**Supplemental Figure S1. *P. aeruginosa* PAO1 cell and supernatant enhance *S. sanguinis* growth.** (A) *S. sanguinis* SK36 was grown in monoculture or in coculture with *P. aeruginosa* PAO1. This finding replicate findings from a previous study from our lab (8). (B) *P. aeruginosa* supernatants also enhance the growth of *S. sanguinis* SK36. *P. aeruginosa* supernatants were prepared by two-fold serial dilutions with fresh MEM+L-Gln medium, and the dilution factors is indicated on the plot. *S. sanguinis* SK36 in monoculture (labeled SK36) and *S. sanguinis* SK36 plus *P. aeruginosa* PAO1 cells (labeled PAO1) in coculture are included as controls. Viability is plotted as $log_{10}$ transformed CFU/ml. Significance was determined by one-way ANOVA followed by Turkey's multiple comparison (**, $P < 0.01$).

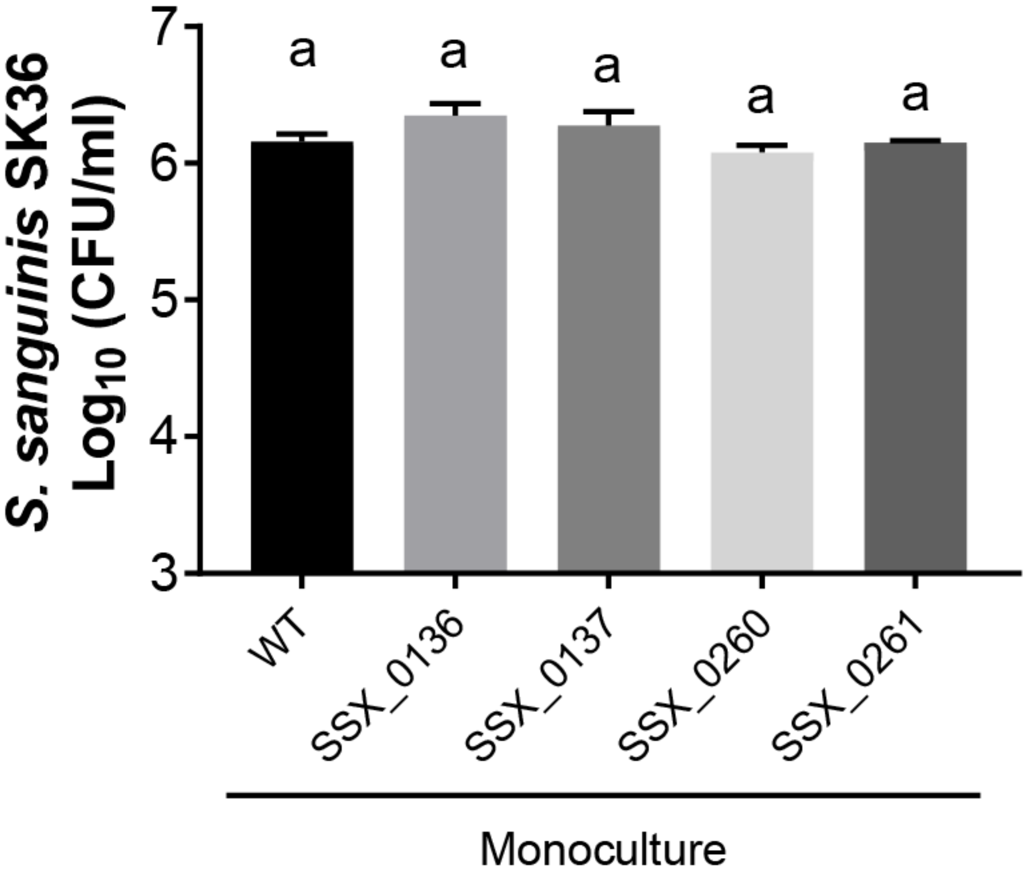


**Supplemental Figure S2. Growth of *S. sanguinis* mutants in monoculture.** Shown is the growth, expressed as $\log_{10}$ (CFU/ml), of the wild type and the indicated *S. sanguinis* mutants in monoculture. a; no significant difference between the WT and any of the mutants.

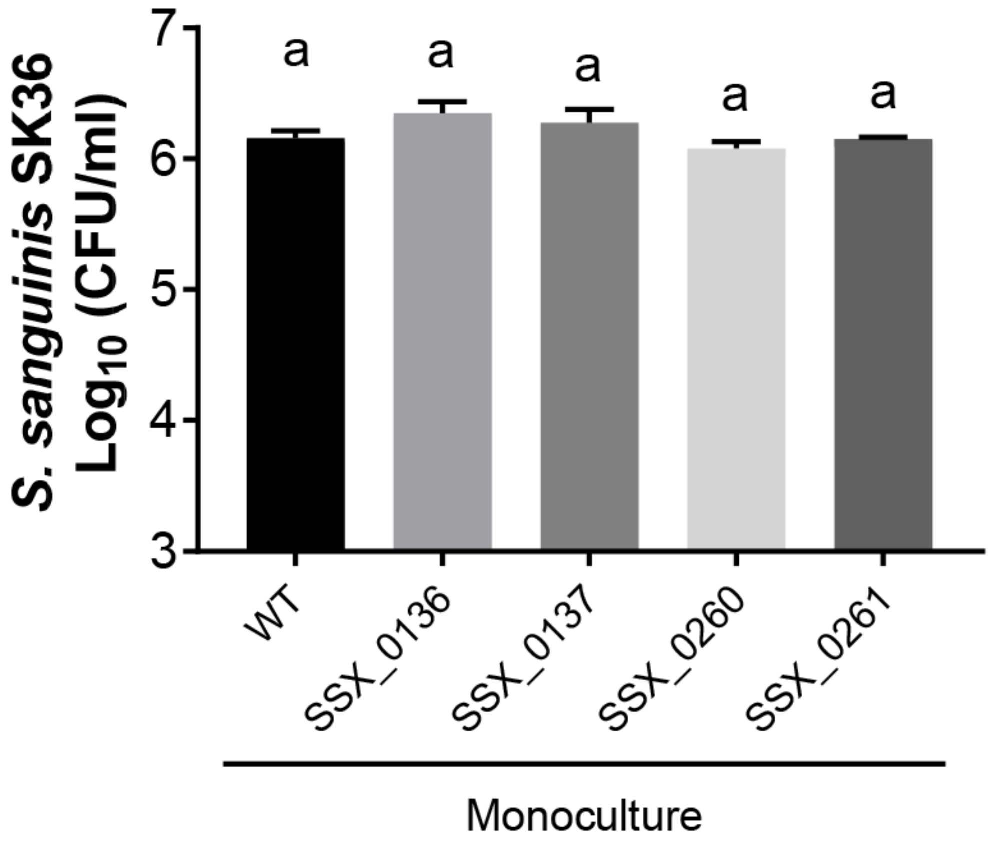


**Supplemental Figure S3. Growth of *P. aeruginosa* in monoculture and coculture with zinc supplementation.** Growth of *P. aeruginosa*, expressed as $\log_{10}$ (CFU/ml), in media with or without addition of zinc at the indicated concentration. a; no significant difference in any of the tested zinc concentrations.

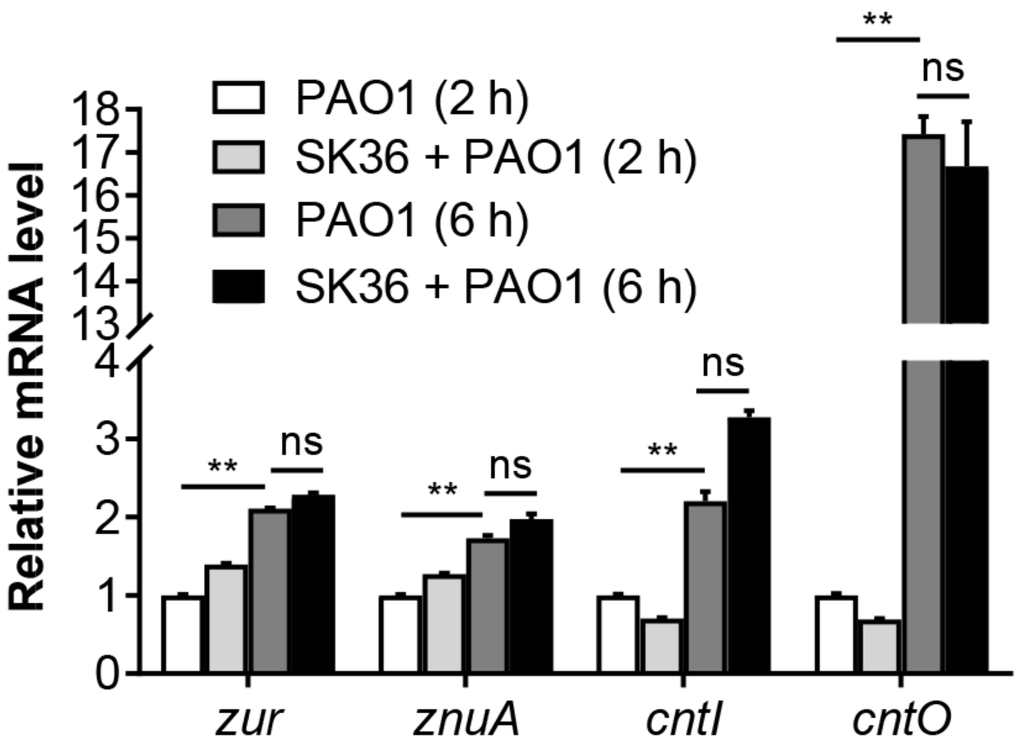


**Supplemental Figure S4. The expression of *P. aeruginosa* zinc uptake and regulator genes are unaffected by the presence of *S. sanguinis*.** Relative mRNA expression of *P. aeruginosa* zinc uptake (*zunA*, *ctnO* and *ctnI*) and regulator (*zur*) genes in monoculture and coculture with *P. aeruginosa* at 2 h and 6 h. The relative mRNA expression was measured using qRT-PCR, normalized to the expression of the *PA2875* control gene, and calculated using the $2^{-\Delta\Delta CT}$ method setting the value of PAO1 (2 h) as one. Significance was determined by ANOVA with Turkey's multiple comparison test (**, $P$ < 0.01).

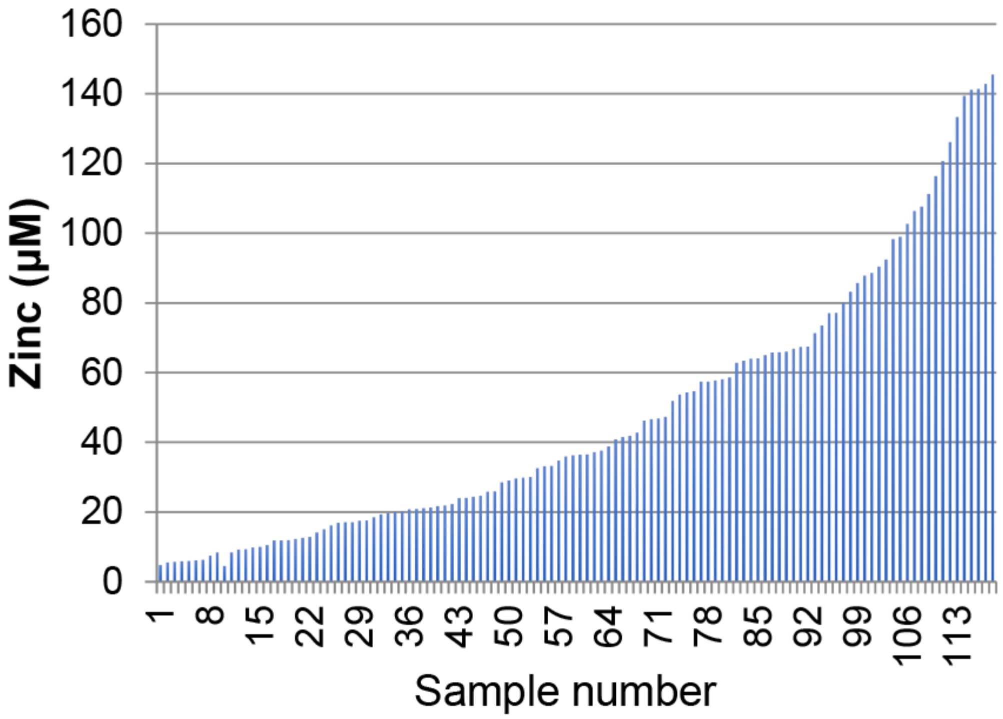


**Supplemental Figure S5. Concentration of zinc in sputum samples.** Shown is the concentration of zinc measured in each of the 118 sputum samples analyzed, sorted from low to high concentration of zinc. The zinc concentration is plotted on the Y axis in μM.
